# Bacteria hyperpolarize in response to oxygen drop

**DOI:** 10.64898/2026.10.07.757406

**Authors:** Maya Peters Kostman, Joshua Girard, Wen Ma, Joseph W. Larkin

## Abstract

Aerobic microbes use oxygen as a terminal electron acceptor to establish an electrochemical gradient across their cell membranes. While this membrane potential (ΔΨ) is essential for multiple cellular processes, it is unknown how it is maintained in bacteria during shifts in oxygen availability. We find that bacteria hyperpolarize in response to oxygen drop. Focusing on *Bacillus subtilis* and guided by modeling, we report that hyperpolarization magnitude scales with electron acceptor deprivation. We find that potassium efflux contributes to the response. Surveying mutants, we uncover genes involved in potassium homeostasis and oxygen stress that contribute to hyperpolarization. We show that hyperpolarization scales with faster growth rates resulting from nutrient type and colony spatial heterogeneity. Finally, we demonstrate that hyperpolarization enhances cellular growth recovery, at the cost of increased antibiotic susceptibility. Our results demonstrate a conserved link between ΔΨ and oxygen stress, suggesting a common mechanism for responding to shifting oxygen availability.

## Introduction

Bacteria rely on membrane potential (ΔΨ), the separation of charges across the membrane, to power crucial processes such as nutrient transport (Lolkema et al., 1998; Simoni & Postma, 1975; Tolner et al., 1995), flagellar motor rotation (Biquet-Bisquert et al., 2021), and oxidative phosphorylation (Harold, 1972). In aerobic bacteria, oxidative phosphorylation provides a major source of energy by flowing protons through the ATP synthase to generate ATP while using oxygen as a final electron acceptor for the electron transport chain (ETC) (Mitchell, 1961). This proton flow is mediated by the proton motive force (PMF), which is primarily composed of the ΔΨ of the cell (Berry et al., 2018; Nicholls & Ferguson, 2013). ΔΨ and oxygen consumption are consequently closely linked.

Oxygen is a highly fluctuating resource in bacterial environments. In the soil, oxygen levels drop following rainfall (Sierra & Renault, 1998), while in the gut, injury can rapidly alter intestinal oxygenation (Zong et al., 2024). ΔΨ has also been shown to be highly dynamic. Ion flux can drive changes in ΔΨ and has been shown to mediate processes such as osmoregulation (Stautz et al., 2021), pH homeostasis (Stautz et al., 2021), mechanosensation (Bruni et al., 2017), and nitrogen stress (Prindle et al., 2015). Bacteria contain several ΔΨ-dependent proteins that control the transport of molecules across the membrane (Kaback, 2015; Tolner et al., 1995). Furthermore, several proteins have been identified to have ΔΨ-dependent localization (Strahl & Hamoen, 2010). Understanding ΔΨ also holds clinical relevance as the uptake of multiple antibiotics has been shown to be ΔΨ-dependent (Bruni & Kralj, 2020; Damper & Epstein, 1981; Ezraty et al., 2013). Changes to ΔΨ can, therefore, have wide ranging consequences for molecular physiology.

The dynamic nature of oxygen conditions and ΔΨ, and their connection raises the question of whether and how aerobic microbes maintain ΔΨ in response to changes in oxygen. Because oxygen powers ΔΨ, it is expected that oxygen deprivation would result in dissipation of ΔΨ. Previous experiments, however, show mixed ΔΨ responses to oxygen depletion. Liquid bacterial cultures responded differently to low oxygen, with some hyperpolarizing and others depolarizing (Kirchhoff & Cypionka, 2017). Arjes, et al. found that *Bacillus subtilis* depolarized in response to oxygen depletion by cellular respiration (Arjes et al., 2020), while Hennes et al. saw low oxygen induced shells of hyperpolarization in colonies of *Neisseria gonorrhoeae* (Hennes et al., 2023). Jin, et al. found that bacterial ion flux was regulated by metabolic state (Jin et al., 2023). No study, however, has systematically investigated the bacterial ΔΨ response to rapid and controlled shifts in oxygen in real time and recorded its physiological consequences.

In this paper, we directly investigate the ΔΨ response to external oxygen shifts in multiple species of aerobic bacteria. Using time-lapse imaging with controlled environmental oxygen concentrations and fluorescent ΔΨ reporters, we find that bacteria hyperpolarize in response to rapid drops in oxygen availability. Cells exhibit a larger magnitude of ΔΨ response when present at higher density or when subjected to a larger applied oxygen drop, demonstrating that hyperpolarization scales with the local degree of oxygen deprivation. To examine the effect of cell density and oxygen consumption on the local oxygen shift experienced by cells and their consequent ΔΨ responses, we create a computational model that predicts cell-specific oxygen uptake based on ambient oxygen availability and colony geometry. The model supports a role for local oxygen competition in determining the magnitude of hyperpolarization. We next identify potassium as an ion involved in mediating the response and show that knocking out genes related to potassium homeostasis as well as oxygen and redox stress hampers hyperpolarization. We hypothesize that hyperpolarization would be stronger for cells with higher oxygen consumption and test this by varying media composition, resulting in different growth rates. We find that the magnitude of hyperpolarization also depends on growth rate, where faster growing cells hyperpolarize more. We show that this pattern also plays out within the nutrient heterogeneity generated in colonies, where faster growing edge cells with more access to nutrients are more hyperpolarized. Finally, we show that hyperpolarization is important for cell proliferation following oxygen drop and find that the bactericidal effect of aminoglycoside antibiotics increases during periods of oxygen shift-induced hyperpolarization. Together, our results show a conserved physiological response that maintains ΔΨ in changing oxygen conditions.

## Results

### Bacteria hyperpolarize during oxygen downshift and depolarize upon oxygen upshift

To determine ΔΨ response to shifts in oxygen, we imaged microcolonies of *B. subtilis* cells on agarose pads with MSgg media (Methods) while undergoing a shift from air (∼21%) to 1% oxygen for 2 hours, before returning to air once again (Supplementary Fig. 1A, Methods). We confirmed that oxygen shifted *in situ* using the oxygen-sensing dye RTDP (Kannan et al., 2023) (Supplementary Fig. 1B, Methods).

We monitored ΔΨ with the fluorescent Nernstian reporter dye thioflavin T (ThT), which is trafficked into or out of cells according to their ΔΨ (Kralj et al., 2011; Strahl & Hamoen, 2010). *B. subtilis* cells showed a stark increase in ThT fluorescence in 1% oxygen, indicating hyperpolarization (Fig. 1A-B, Movie 1). ThT signal peaked after 1 hour in low oxygen. Fluorescence decreased back sharply to levels similar to before the shift within 30min of oxygen being restored, indicating depolarization. Hyperpolarization during oxygen downshift was also observed using the ΔΨ dyes Tetramethylrhodamine methyl ester (TMRM) (Kralj et al., 2011) (Supplementary Fig. 1E-F) and 3,3’-Dipropylthiadicarbocyanine Iodide (DISC3(5)) (Strahl & Hamoen, 2010) (Supplementary Fig. 1G-H).

**Figure 1.**
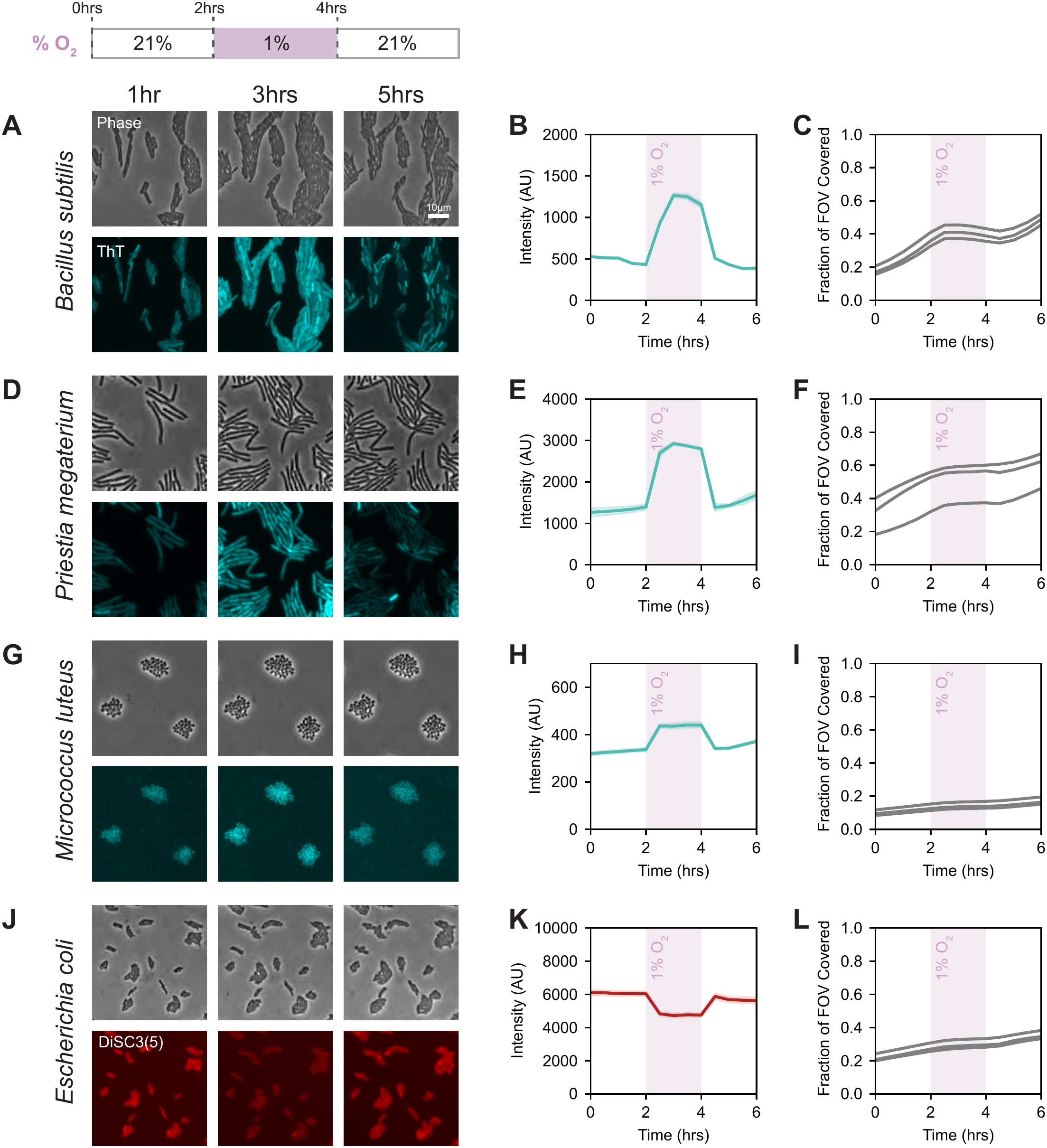
Bacteria hyperpolarize during oxygen downshift and depolarize upon oxygen upshift. Oxygen was dropped from air (21%) to 1% between 2-4 hrs for *B. subtilis* (**A-C**), *P. megaterium* (**D-F**), *M. luteus* (**G-I**), or *E. coli* (**J-L**). **A, D, G, J.** still images of cells at 1hr (left) 3hrs (middle) and 5hrs (right) in phase (top) and fluorescent membrane potential reporter ThT (bottom). In **J**, DISC3(5) was used in place of ThT for better absorption by Gram-negative *E. coli*. Note that DISC3(5) behaves inversely to ThT, with a decrease in fluorescence indicating hyperpolarization. **B, E, H, K.** Mean fluorescence intensity over time for either ThT (**B, E, H**) or DISC3(5) (**K**). N = 3 fields of view. Shaded region indicates mean +/- s. e. m. **C, F, I, L.** fFOV (fraction of the field of view covered by cell monolayer) over time for 3 fields of view measured. Pink blocks denote the duration of 1% oxygen.

As expected, cells slowed growth during periods of low oxygen such that the increase the fraction of the field of view covered by cell monolayer (fFOV, taken as a measurement of 2D cell density), plateaued during low oxygen conditions, coinciding with hyperpolarization (Fig. 1C, Supplementary Fig. 1J, Movie 2). Growth resumed after oxygen was restored (Fig. 1C).

We next asked if other bacterial species exhibited a similar oxygen-induced ΔΨ response. We observed the same membrane polarization pattern for two other species of Gram-positive bacteria: *Priestia megaterium* and *Micrococcus luteus*, as well as the Gram-negative *Escherichia coli* (Fig. 1D-L), suggesting that the physiological response of hyperpolarization is exhibited by a range of aerobic organisms.

These results demonstrate that, across broad phyla, bacteria respond to transient oxygen drops by rapidly hyperpolarizing their ΔΨ. We next investigated what factors determined the degree of hyperpolarization.

### Hyperpolarization magnitude scales with the degree of electron acceptor deprivation

Having established oxygen-induced ΔΨ dynamics across several species, we next focused on *B. subtilis* to investigate the behavior in more detail. We observed that for the same drop in supplied oxygen, *B. subtilis* cells hyperpolarized more if they were present at higher density and fFOV (Fig. 2A-B). We hypothesized that this was due to the depletion of oxygen by neighboring cells, which reduced local oxygen availability. To disentangle the effects of oxygen diffusion, cell density, and cellular hyperpolarization response, we developed a steady-state reaction-diffusion model that explicitly represents individual cells and predicts oxygen uptake as a function of ambient oxygen availability and colony geometry (Methods, Supplementary Information). In the model, we populate a field of view with cells at a specified average density and external oxygen concentration, which allows us to separate the two factors. Each cell takes up oxygen at a rate proportional to its local oxygen concentration and hyperpolarizes if this rate dips below a threshold. As cells consume dissolved oxygen, which diffuses at a limited rate, they create a local oxygen-deprivation well. At high density, overlap between local depletion regions causes cells to be more oxygen-starved locally (Fig. 2C). The model predicts that cellular hyperpolarization depends on both external oxygen drop and cell density (Fig. 2D). Both stronger oxygen drops and higher cell densities are predicted to result in lower local oxygen availability and, consequently, greater hyperpolarization. Furthermore, because the influence of each cellular deprivation well is spatially limited and cells respond only once their uptake falls below the threshold, we see that in sufficiently oxygen-rich environments, cells remain unstressed and do not hyperpolarize until a density threshold is achieved. Overall, ΔΨ response is predicted to depend on three major factors: the ambient oxygen supplied to the chamber, the density of surrounding cells competing for oxygen, and the oxygen demand of the cells themselves.

**Figure 2.**
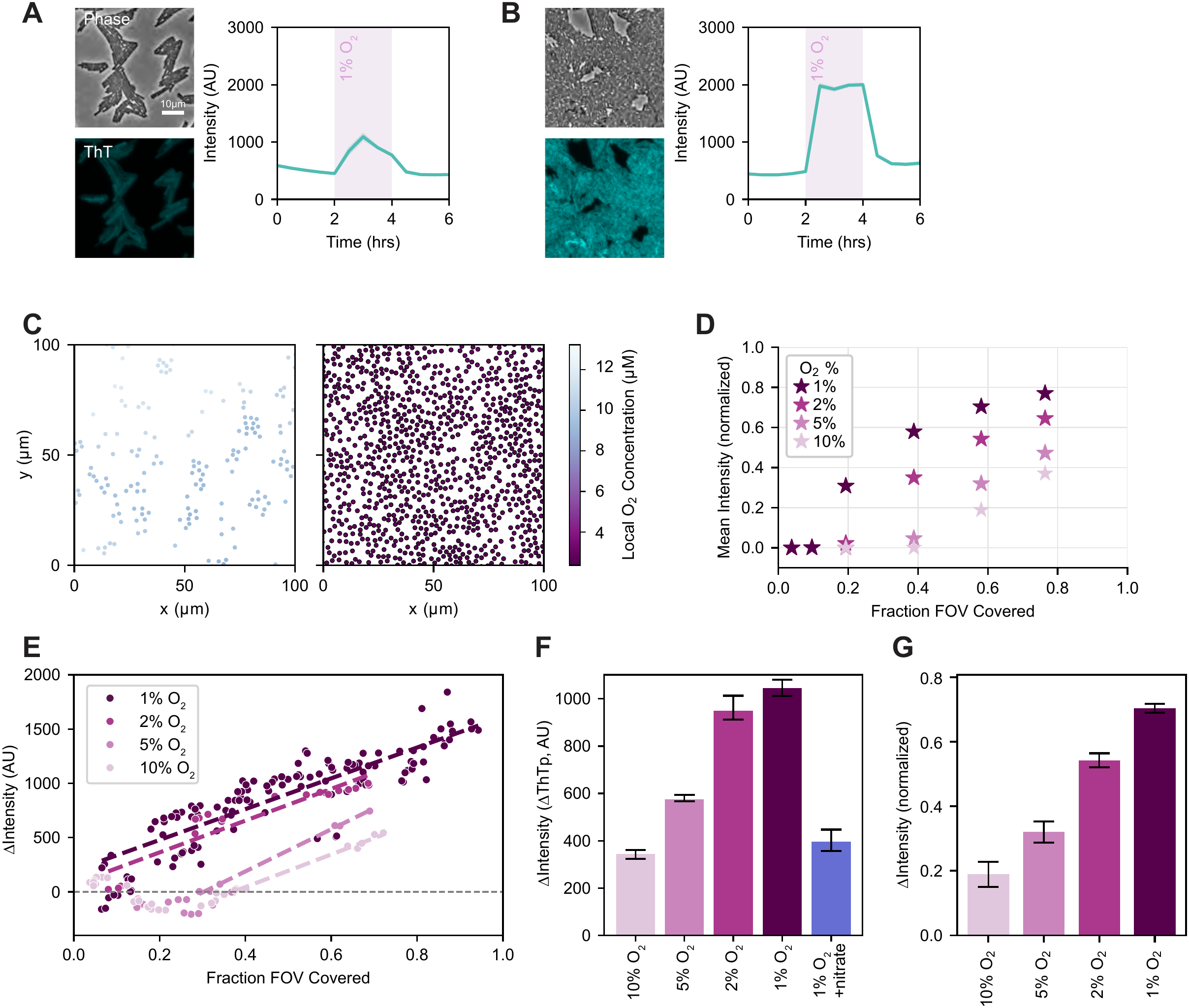
Hyperpolarization magnitude scales with the degree oxygen deprivation. Representative images for *B. subtilis* at 3 hrs at low cell density (low fFOV) (**A**) and high cell density (high fFOV) (**B**) in phase (top left) and ThT (bottom left). Graphs show mean ThT fluorescence intensity over time (right). N = 3 fields of view. Shaded region indicates mean +/- s. e. m. Pink blocks denote the duration of 1% oxygen. **C.** Simulated *B. subtilis* colonies of low density (.09 fFOV, left) and high density (.58 fFOV, right) after oxygen downshift to 2%. Color represents numerically evaluated local oxygen level. **D.** Scatterplot for simulated change in mean ThT intensity at steady state versus fFOV, using parameters fitted to the data in **E** (Methods). Colors correspond to oxygen level after downshift. Each star represents one evaluation of a simulated colony as shown in (**C**). **E.** Experimental scatterplot for change in ThT intensity after 1 hr of oxygen downshift to either 1%, 2%, 5% or 10% oxygen versus fFOV at the time of oxygen downshift. Each point represents one field of view imaged. Dashed lines represent linear regression for points with positive net intensity and to the right of the first negative Δintensity value. N ≥ 6 for each condition. **F.** Change in intensity predicted by linear regression from (**E**) at .6 fFOV, as well as for a condition including nitrate (blue bar). Error bars represent the 95% confidence interval for the linear regression at .6 fFOV. (N ≥ 6 for each condition) **G.** Change in normalized intensity determined from simulation evaluated at .6 fFOV. Error bars represent +/- 1 std. dev. of the intensity distribution within the simulated population.

To determine if cells responded to oxygen shifts in a density-dependent manner as predicted by the model, we performed imaging experiments with varied levels of both supplied oxygen percentage and cell density. We manipulated external oxygen by shifting our input from air to 10%, 5%, 2% or 1% oxygen and we generated different fFOVs by inoculating different concentrations of cells on the imaging pad (Methods). To determine the strength of the ΔΨ response, we measured the net change in mean ThT fluorescence after 1 hour of low oxygen (ΔThT). Consistent with our model predictions, we found that ΔThT increased with fFOV, indicating that denser communities hyperpolarized more (Fig. 2E). ΔThT also increased with lower oxygen percentages, further confirming the response to be dependent on oxygen availability. For the weaker oxygen drops, 10% and 5%, we saw that cells only hyperpolarized once they reached a high enough density, consistent with our model predictions of a cell density threshold for hyperpolarization in more oxygen-rich conditions. To quantify and directly compare the effects of different oxygen drops on hyperpolarization, we performed linear regression and compared ΔThT at .6 fFOV, which we refer to as ΔThTp (Fig. 2F). ΔThTp showed the oxygen dependence of hyperpolarization is consistent with the model (Fig. 2G).

We next asked if the impact of oxygen on ΔΨ was specific to its role as an electron acceptor. It is known that *B. subtilis* can use nitrate as an alternative electron acceptor (Bueno et al., 2012). We hypothesized that supplying cells with nitrate would compensate for the loss of electron acceptor in low oxygen and reduce hyperpolarization. We supplemented media with nitrate and saw that cells subjected to a 1% oxygen shift had lower ΔThTp than in the 1% shift without nitrate, suggesting that this condition was instead more similar to the 10% oxygen shift condition and that NaNO_3_ could compensate to some extent for the lack of oxygen (Fig. 2F). These results demonstrate that the degree of cell hyperpolarization in response to oxygen scales with the degree to which cells are deprived of a terminal electron acceptor. We next asked what mechanisms were involved in cellular transduction of oxygen shifts into membrane hyperpolarization.

### Potassium efflux contributes to hyperpolarization

Previous work has shown that potassium efflux can mediate transient membrane hyperpolarization during nitrogen stress in *B. subtilis* (Prindle et al., 2015). To determine whether potassium flux occurs during oxygen shift, we measured the amount of potassium in the environment using the membrane-impermeable extracellular potassium indicator IPG-4. During downshifts to 10% oxygen, cells showed an increase in the rate of extracellular potassium accumulation, which decreased once oxygen was restored (Fig. 3A, Supplementary Fig. 3A). This shows that potassium release from cells increases during the oxygen drop, which would contribute to cell hyperpolarization. Potassium efflux is known to cause a decrease in cell turgor pressure, which results in changes to cell volume (Brogan et al., 2025; Commichau et al., 2018). We measured the mean single cell area for cells undergoing a drop to 1% oxygen and found that cells decreased in size during low oxygen (Fig. 3B), consistent with potassium efflux. We also confirmed this to be the case in the absence of ThT dye (Supplementary Fig. 3D). We measured the average change in single cell area following 1 hour of 1% oxygen and found that change in cell area scaled linearly with change in ΔΨ such that cells that hyperpolarized more also shrunk more (Fig. 3C). This suggests that stronger hyperpolarization may coincide with higher potassium release.

**Figure 3.**
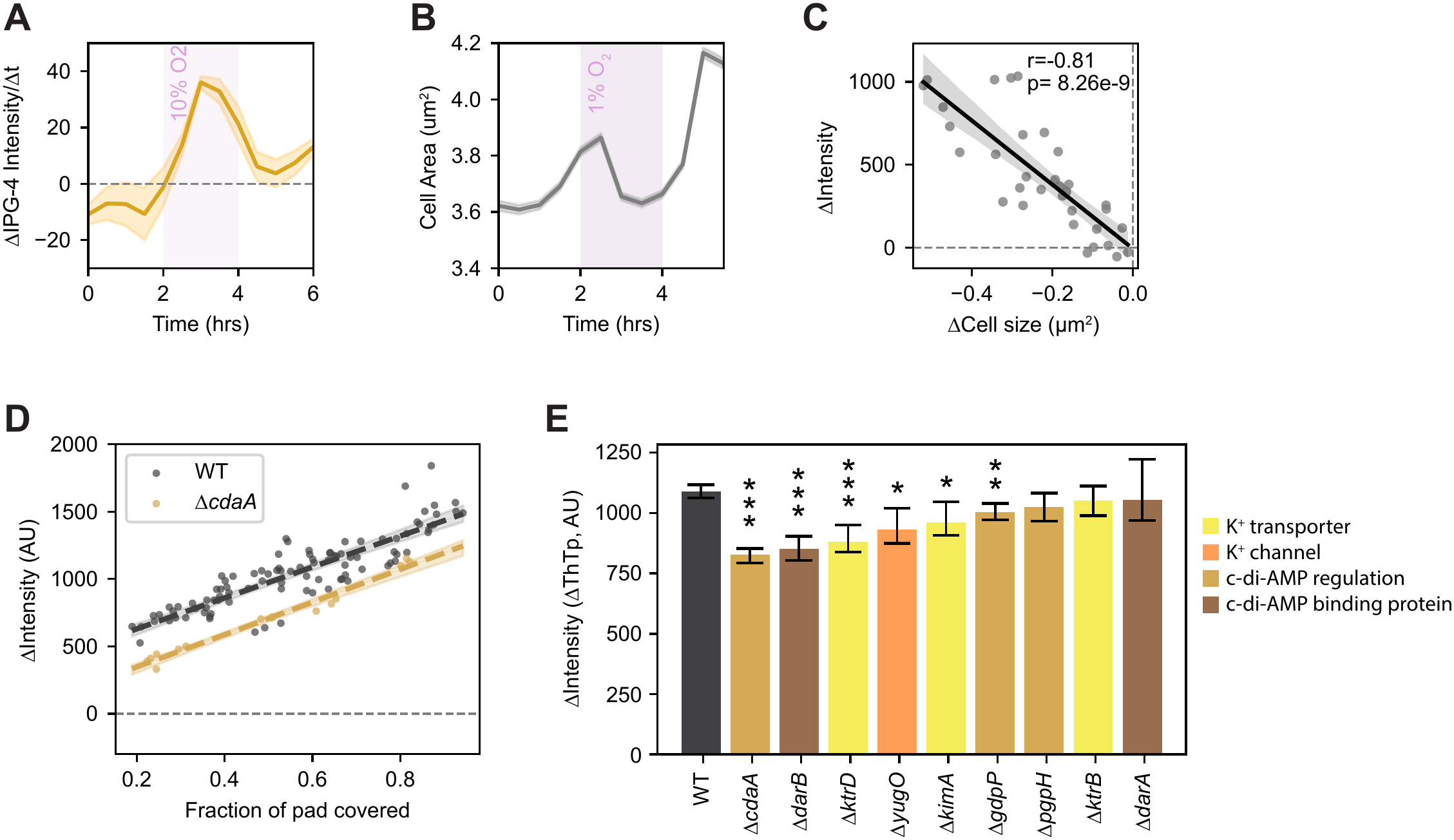
Potassium efflux contributes to hyperpolarization. **A.** Mean change in IPG-4 extracellular potassium dye fluorescence intensity over time. N = 3 fields of view. Shaded region indicates mean +/- s. e. m. Pink block denotes the duration of 10% oxygen. **B.** Mean single cell area over time. N = 3 FOVs. Shaded region indicates mean +/- s. e. m. Pink block indicates the duration of 1% oxygen. **C.** Scatterplot for change in ThT intensity versus the change in single cell area after 1 hr in 1% oxygen. **D.** Scatterplot for change in ThT intensity after 1 hour of downshift to 1% oxygen versus fFOV at the time of oxygen downshift for WT (grey) and Δ*cdaA* (yellow). Each point represents one field of view imaged. Lines represent linear regression and shaded region shows the 95% confidence interval for the regression. **E.** Predicted change in ThT intensity after 1 hour of 1% oxygen at .6 fFOV derived from linear regression, as seen in **D**, for different gene knockouts (N ≥ 5 for each knockout). Error bars represent the 95% confidence interval for the linear regression at .6 fFOV. Stars indicate significance (* p value <.05, ** p value <.01, *** p value <.001).

Given our observation of potassium efflux during hyperpolarization, we next asked whether proteins involved in potassium transport impacted the response. We imaged 9 different knockout strains for genes involved in potassium homeostasis and measured ΔThT, fFOV and ΔThTp as described above. Several mutants showed decreased ΔThTp compared to WT (Fig. 3D-E). These included the potassium transporters *ktrD* and *kimA* and the potassium channel *yugO* (Stautz et al., 2021). We also saw a decrease in ΔThTp for genes involved in regulating the second messenger c-di-AMP, which controls potassium homeostasis by blocking the expression and activity of potassium transporters (Yin et al., 2020). c-di-AMP is synthesized by diadenylate cyclases and broken down by phosphodiesterases (Yin et al., 2020). The knockout for the main diadenylate cyclase involved in potassium homeostasis, *cdaA* (Gundlach, Herzberg, Kaever, et al., 2017), showed the strongest decrease in ΔThTp. One of the phosphodiesterases, *gdpP* but not *pgpH*, also showed slightly reduced ΔThTp. We also saw a significant decrease in ΔThTp for the c-di-AMP binding protein knockout *darB,* which activates the stringent response when c-di-AMP is low (Krüger et al., 2021). These results highlight the role of potassium flux and homeostasis as well as downstream pathways in defining hyperpolarization during oxygen downshift.

### Oxygen stress genes impact hyperpolarization response to oxygen downshift

Oxygen shift-mediated hyperpolarization connects to many aspects of cellular physiology, including oxygen sensing, metabolism, and redox state. We hypothesized that genes involved in these pathways could play a role in inducing hyperpolarization. We therefore generated single knockout strains of 17 genes involved in either oxygen, redox or metabolic stress, as well as the ETC (Gray et al., 2024; Härtig & Jahn, 2012). We compared their ΔThTp during shifts from air to 1% oxygen as described above (Fig. 4A-B). All tested strains hyperpolarized in response to the oxygen drop, but 3 mutants showed drastically attenuated hyperpolarization (Fig. 4B). These were all involved in redox and oxygen sensing: Δ*ndh*, Δ*rex* and Δ*resD*. *ndh* encodes the main NADH dehydrogenase in *B. subtilis*, responsible for converting NADH into NAD+ and feeding electrons into the ETC (Gyan et al., 2006; Härtig & Jahn, 2012). *rex* encodes the redox-sensing transcription factor Rex, responsible for activation of fermentation genes in response to elevated NADH/NAD+ ratio (Härtig & Jahn, 2012). *resD* encodes part of the two-component system ResDE that senses oxygen availability (Härtig & Jahn, 2012). ResE is the membrane component and phosphorylates ResD in conditions of low oxygen. Once activated, ResD dissociates from the membrane and activates 27 genes and operons associated with anaerobic respiration including *fnr*, *yclJK*, and *ldhlctP* (Härtig & Jahn, 2012). Δ*resE* also showed significant reduction in hyperpolarization, although not as strongly as Δ*resD*, which is consistent with studies that have seen that ResD is able to respond to redox and oxygen stress even in the absence of phosphorylation by ResE (Geng et al., 2004).

**Figure 4.**
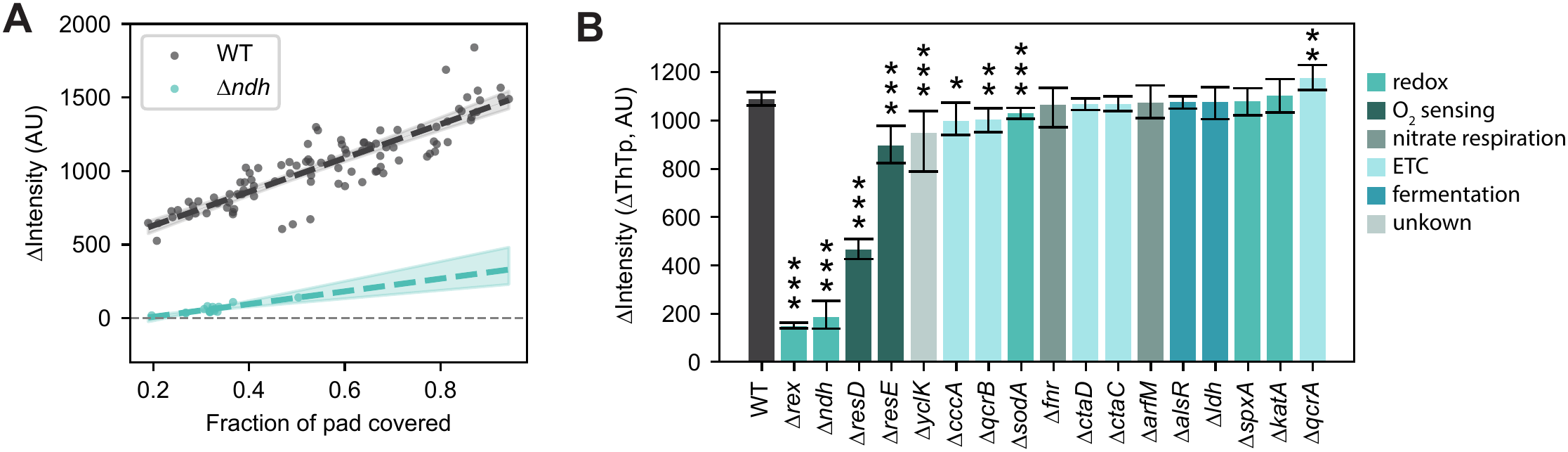
Oxygen stress genes impact hyperpolarization response to oxygen downshift. **A.** Scatterplot for change in ThT intensity after 1 hour of oxygen downshift to 1% oxygen versus fFOV at the time of oxygen downshift for WT (grey) and Δ*ndh* (cyan). Each point represents one field of view imaged. Lines represent linear regression and shaded region shows the 95% confidence interval for the regression. **B.** Predicted change in ThT intensity after 1 hour of 1% oxygen downshift at .6 fFOV derived from linear regression, as seen in A, for different gene knockouts (N ≥ 4 for each knockout). Error bars represent the 95% confidence interval for the linear regression at .6 fFOV.

We also measured ΔThTp for 3 genes regulated by *resD* at low oxygen: the oxygen regulator gene *fnr*, the two-component system *yclK*, and the lactate dehydrogenase *ldh* (Härtig & Jahn, 2012). Of these, only *yclK* showed a significant reduction in hyperpolarization. The gene *yclK* encodes a potential sensor kinase that composes part of the YclJK two component system, which is upregulated by ResD in low oxygen (Härtig et al., 2004; Härtig & Jahn, 2012). The activating signal and regulated downstream genes for YclJK are still unknown, but our results indicate it could play a role in mediating hyperpolarization. Δ*qcrA* was the only gene knockout that resulted in a slightly increased hyperpolarization response to oxygen drop. QcrA is the iron-sulfur subunit of respiratory complex III and has been shown to have ΔΨ-dependent localization (Gray et al., 2024). Altogether, our survey of knockout strains revealed oxygen shift-mediated hyperpolarization is a response connected to many aspects of cellular physiology, including oxygen sensing, metabolism, and redox state.

Our knockout strains also showed a correlation between hyperpolarization and growth rate, where strains with longer doubling times showed less change in ThT intensity (Supplementary Fig. 4B). This raised the possibility that growth rate and metabolism could themselves be determinants of hyperpolarization.

### Magnitude of hyperpolarization scales with growth rate in both microcolonies and biofilms

Our model predicts that cellular oxygen demand is one of the determinants of hyperpolarization magnitude. This prediction suggests that faster growing cells may require more oxygen and hyperpolarize more during oxygen drops. This trend is consistent with our gene knockout data (Supplementary Fig. 4B). To assess the relationship between hyperpolarization and growth rate, we performed experiments with wildtype cells in nutrient sources with varied doubling times or with antibiotics that slow growth (Fig. 5A, Supplementary Table 2). These conditions also resulted in a range of ΔThTp values (Fig. 5B). We saw a negative correlation between doubling time and hyperpolarization: conditions with faster growth (i.e. lower doubling times) resulted in a greater ThT change in response to oxygen drops (Fig. 5C). This is in line with our model where slower-growing cells with a lower oxygen demand experience less oxygen deprivation and hyperpolarize less.

**Figure 5.**
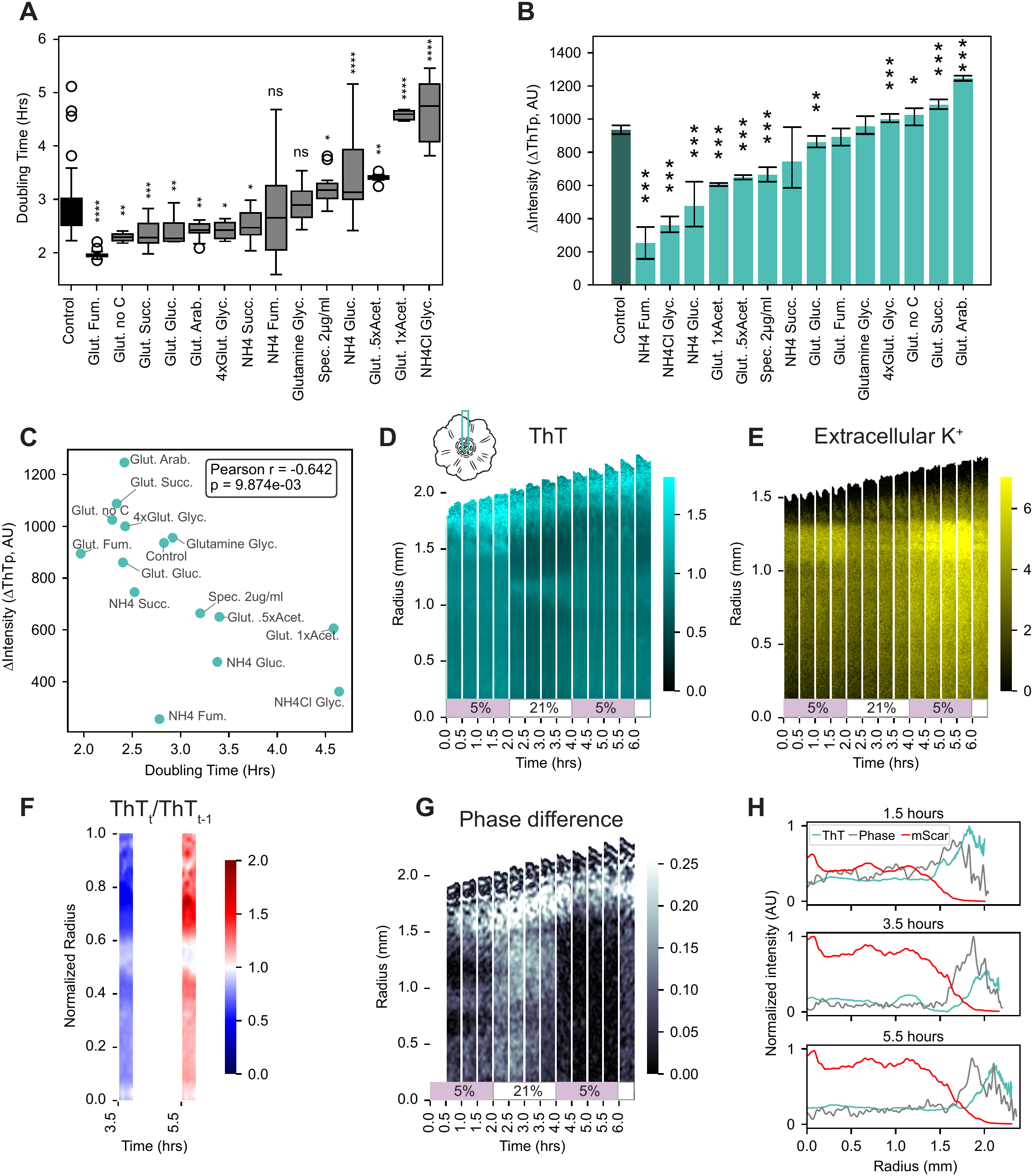
Magnitude of hyperpolarization scales with growth rate in both microcolonies and biofilms. **A.** Mean doubling time in 21% oxygen (air) for WT *B. subtilis* grown on different nutrient sources or antibiotic. See Supplementary Table 2 for condition descriptions. Error bars represent s. e. m. N ≥ 2 for each condition. **B.** Predicted change in ThT intensity after 1 hr of 1% oxygen downshift at .6 fFOV derived from linear regression for WT *B. subtilis* grown on different nutrients or antibiotic. Error bars represent the 95% confidence interval for the linear regression at .6 fFOV. N ≥ 2 for each condition. Stars indicate significance (* p value <.05, ** p value <.01, *** p value <.001). **C.** Scatterplot for ΔThT (shown in B) versus the average doubling time for each condition (shown in A). Pearson r shows the Pearson correlation and p shows the p-value. **D, E.** Radial strips (D inset) of 18 hr-old biofilms undergoing oxygen shifts from 5% oxygen to 21% (air) every 2 hours shown in normalized ThT fluorescence (D) and IPG-4 extracellular potassium dye fluorescence (E). Colorbars represent normalized fluorescence and raw fluorescence respectively. **F.** Normalized ThT fluorescence from D at 3.5 hrs divided by fluorescence at 1.5 hrs (switch from 5% to 21% O2) and normalized ThT fluorescence at 5.5 hrs divided by fluorescence at 3.5 hrs (switch from 21% to 5% O2). Radial coordinates have been scaled by the colony radius. **G.** Radial strips from the colony in D shown in the phase difference calculation. Color represents change in phase signal. **H.** Normalized intensities for normalized ThT, phase difference, and mScarlet across the radius of the colony shown in (**D**) and (**G**) at 3 different time points.

We expected that the same relationship between growth rate and hyperpolarization might also play out within a heterogeneous community. Bacteria are commonly found in biofilms, where nutrient conditions fluctuate dramatically over the colony and cells at the edge have better access to nutrients (Liu et al., 2015; Mulder et al., 2024). We therefore hypothesized that ΔΨ responses to changing oxygen would vary spatially throughout the colony, with nutrient-rich exterior cells exhibiting a stronger response than starved interior cells. We grew biofilms for 18 hours in the presence of ThT and fluctuated the oxygen every 2 hours between air and 5% oxygen. We observed that, as seen in previous experiments, the total community ThT signal (normalized by constitutively expressed mScarlet) increased during low oxygen intervals (Fig. 5D, Movie 3), accompanied by an increase in extracellular potassium (Fig. 5E). We saw a stark boundary in hyperpolarization emerge where the edge of the biofilm was consistently more hyperpolarized than the center regardless of oxygen level supplied, and the edge also had the most temporal change in ThT signal (Fig. 5F). It has been shown that biofilm growth occurs mainly at the edge (Jones et al., 2026; Liu et al., 2015), a behavior that can be seen in our measurements of phase difference images (Fig. 5G, Movie 4). The region of higher growth coincides with the region of highest ThT signal (Fig. 5H). These results demonstrate that, in dense colonies, nutrient gradients and metabolic heterogeneity cause genetically identical cells to exhibit different responses to oxygen shifts across the colony in a growth rate-dependent manner.

### Hyperpolarization supports cell proliferation following oxygen drop

Having characterized the hyperpolarization response of cells undergoing oxygen downshift, we next asked what consequences this behavior has for cell physiology. Because the response is conserved across distantly related bacteria, we predicted it would have a beneficial impact in surviving oxygen drops and that disrupting hyperpolarization would negatively impact cell proliferation. In order to disturb hyperpolarization, we grew cells on agarose pads containing the potassium ionophore valinomycin, which has been shown to inhibit hyperpolarization via potassium flux in bacteria (Prindle et al., 2015). We saw that valinomycin significantly decreased ΔThTp as compared to DMSO control without changing the doubling time (Fig. 6A, Supplementary Fig. 6A-B). We next measured the instantaneous growth rate as the maximum velocity of fFOV expansion (Vmax). Before oxygen downshift, we saw that valinomycin Vmax was similar to DMSO control (Fig. 6B). However, after oxygen was restored, valinomycin-treated cells took longer to resume growth (Fig. 6C-D). Valinomycin also caused a more negative Vmax, indicating more cell death (Fig. 6D). Vmax remained negative even 1 hour after oxygen was recovered to 21% (air). Meanwhile, DMSO control showed a less negative Vmax and only remained negative for 30min following oxygen recovery. ATP level did not appear different between conditions (Supplementary Fig. 6C). Our results indicate that disturbing hyperpolarization using valinomycin disturbs cells’ ability to recover and proliferate following oxygen downshift.

**Figure 6.**
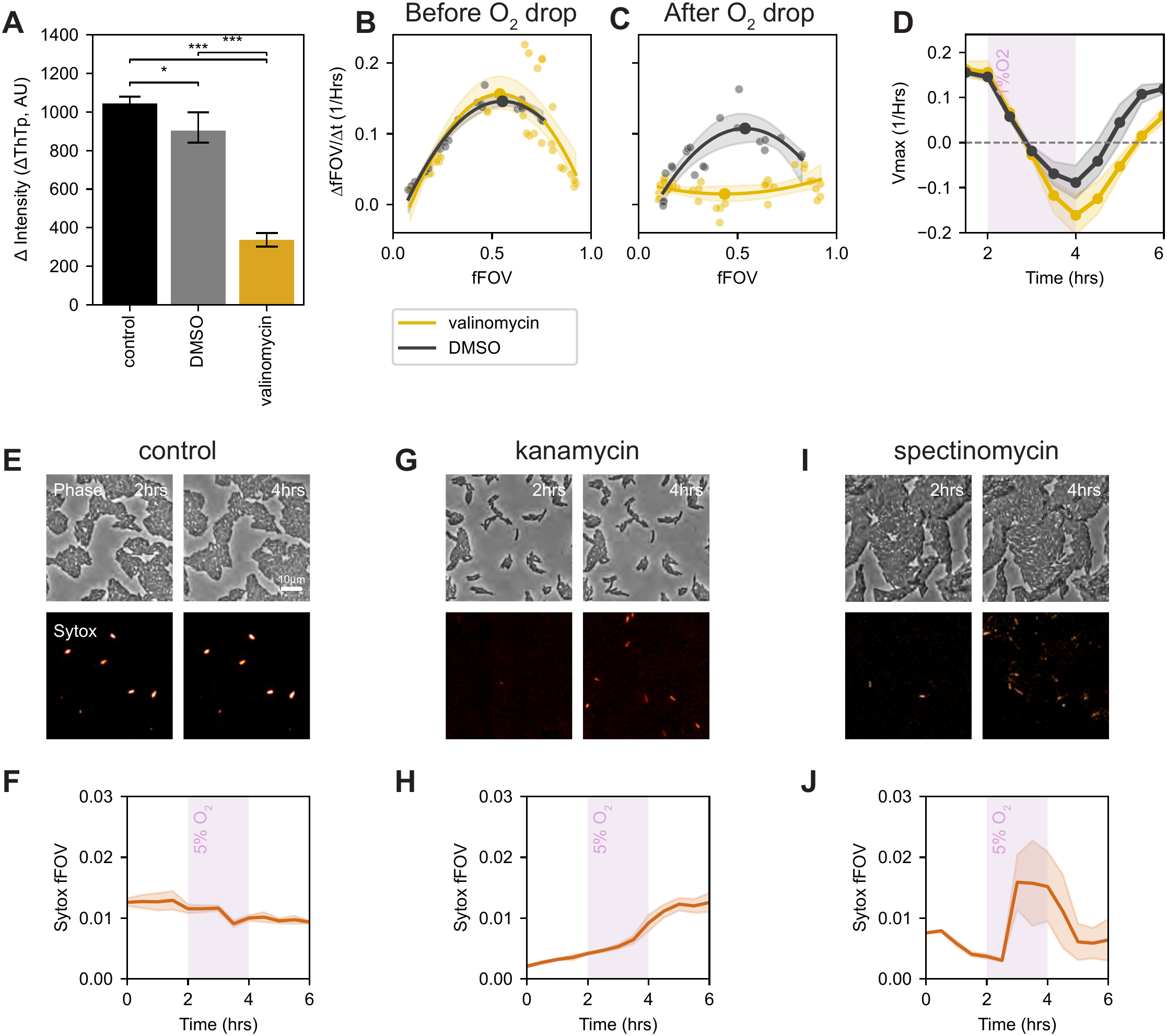
Hyperpolarization supports cell proliferation and increases antibiotic susceptibility. **A.** Change in ThT intensity predicted by linear regression from Supplementary Fig. 6A at .6 fFOV. Error bars represent the 95% confidence interval for the linear regression at .6 fFOV. N ≥ 6 for each condition. **B**, **C**. Rate of change of fFOV (colony expansion rate) for different fFOV’s (fraction of the field of view covered by cell monolayer) before oxygen drop (at 2 hrs) (B), and after oxygen drop (at 5.5 hrs) (**C**). Solid lines represent quadratic regression and the shaded region around the line shows the 95% confidence interval. Solid circles represent the parabola vertex. Yellow represents valinomycin and grey represents DMSO control. **D**. Vertices of the data shown in (B) and (C) plotted for all time points. Shaded region around the line shows the 95% confidence interval. Yellow represents valinomycin and grey represents DMSO control. **E-J**. Oxygen was dropped from air (21%) to 5% between 2-4 hrs for *B. subtilis* in no antibiotic (**E-F**), kanamycin (**G-H**), or spectinomycin (**I-J**). **E, G, I.** still images of cells at 2 hrs (left) and 4 hrs (right) in phase (top) and Sytox Orange cell death stain (bottom). **F, H, J.** Mean fraction of field of view covered by Sytox signal (Sytox fFOV) over time. N = 3 fields of view. Shaded region indicates mean +/- s. e. m.

### Increased bactericidal activity of antibiotics during oxygen downshift

While hyperpolarization allows cell communities to better survive oxygen downshift, we asked whether this behavior could create vulnerability to other environmental challenges. Of particular interest is the impact hyperpolarization has on antibiotic susceptibility. It has been shown that the uptake and activity of aminoglycoside antibiotics relies on ΔΨ (Bruni & Kralj, 2020; Damper & Epstein, 1981; Ezraty et al., 2013). Because cells hyperpolarize in response to oxygen drop, we asked if oxygen downshift increases the effectiveness of the aminoglycoside kanamycin. We grew cells with or without kanamycin in the presence of Sytox to stain for cell death (Choi et al., 2017; Scanlon et al., 2014). We saw an increase in Sytox signal during oxygen drop, which was not present in the control (Fig 6E-H). We also tested the effect of spectinomycin, an aminocyclitol, which is closely related to aminoglycosides. We again saw an increase in cell death during low oxygen (Fig. 6I-J). Our results show that the bactericidal effects of both antibiotics were potentiated during low oxygen, possibly due to the effect of hyperpolarization.

## Discussion

We have shown that bacteria hyperpolarize in response to oxygen downshift and depolarize once oxygen is restored. This behavior is widespread, as it was observed in the Gram-positives *B. subtilis*, *P. megaterium* and *M. luteus*, as well as the Gram-negative *E. coli*. All species tested are able to grow aerobically but show different levels of tolerance to oxygen deprivation, with *M. luteus* considered a strict aerobe (Young et al., 2010), while *E. coli* is a facultative anaerobe known to tolerate oxygen poor environments such as the gut (Martinson & Walk, 2020). Nonetheless, the same pattern of behavior was seen for all species.

We predicted that hyperpolarization would scale with the level of oxygen deprivation experienced locally. We generated a computational model to simulate the physical constraints governing the diffusion and availability of oxygen between cells. Our model suggests that hyperpolarization occurs once cellular oxygen uptake falls below a response threshold and it depends on 1) the global availability of oxygen, 2) the cell density, and 3) the oxygen demand of the cells. These predictions are consistent with our experimental data, where lower supplied oxygen and higher cell density both lead to increased hyperpolarization. In our model, cells with a higher oxygen demand, represented by a higher response threshold, hyperpolarize more at the same oxygen level, which is also consistent with our experimental finding that hyperpolarization scales with growth rate. Faster growing cells, which likely have a higher oxygen requirement, hyperpolarized more strongly. Stratford et al. reported a similar relationship between proliferation and membrane potential for cells subjected to electrical stimulation (Stratford et al., 2019). Our model predictions also allow us to examine quantities that are otherwise difficult to measure, such as per-cell uptake and local oxygen availability, and to extrapolate to untested conditions. The effect of cell density is predicted to result from ‘screening’ behavior, in which denser communities reduce oxygen availability locally by consuming diffusing oxygen and creating depletion zones. Accordingly, hyperpolarization is predicted to depend not just on average cell density, but on spatial arrangement, with more clumped populations potentially exhibiting a stronger response than evenly distributed cells.

Our study revealed that potassium efflux coincides with cell hyperpolarization in both cell monolayers and biofilms, indicating that potassium contributes to hyperpolarization. The involvement of other ions, however, has not been ruled out. Potassium flux is known to regulate turgor pressure (Brogan et al., 2025; Commichau et al., 2018). We see cell shrinkage during low oxygen, which is exacerbated with increasing hyperpolarization, suggesting efflux scales with hyperpolarization. Our mutant experiments showed hindered hyperpolarization for knockouts of *yugO*, *kimA*, and *ktrD*. No knockout completely eliminated hyperpolarization, suggesting redundancy in the system. *yugO* encodes a metabolically gated potassium channel that has been shown to respond to NADH and mediate hyperpolarization during nitrogen stress (Cao et al., 2013; Prindle et al., 2015; Roosild et al., 2002; Schlosser et al., 1993). ΔyugO has previously been reported to attenuate membrane hyperpolarization during electrical stimulation (Stratford et al., 2019). *kimA* encodes a high-affinity proton and potassium symporter in *B. subtilis* (Gundlach, Herzberg, Kaever, et al., 2017; Tascón et al., 2020). Meanwhile, *ktrD* encodes the membrane-spanning channel component of the low-affinity KtrCD potassium transporter, which is thought to act as the main transporter when potassium is abundant (Gundlach, Herzberg, Hertel, et al., 2017). Ktr transporters contain RCK/KTN domains, which have been shown to require ATP to be activated (Teixeira-Duarte et al., 2019) and to change conformation when bound to NADH or NAD+ (Roosild et al., 2002). KtrC also binds ATP and c-di-AMP to change conformation (Rocha et al., 2019). Our mutant experiments also showed hindered hyperpolarization for knockouts of *cdaA*, *gdpP*, and *darB*. *cdaA* and *gdpP* control levels of c-di-AMP, the second messenger regulating potassium homeostasis (Yin et al., 2020). c-di-AMP can signal downstream to activate the stringent response through DarB (Krüger et al., 2021). Therefore, potassium flux seen during low oxygen may also connect to metabolism through c-di-AMP signaling. It is well-documented that acute hypoxia impacts potassium channel activity and expression in eukaryotes (Shimoda & Polak, 2011). Nerve cells have been shown to transiently hyperpolarize which in some cell types is mediated by ATP-sensitive potassium channels (Trapp & Ballanyi, 1995). Our results indicate that the importance of potassium in mediating oxygen shift-induced hyperpolarization extends to bacteria.

Our mutant experiments also revealed a possible role for redox state in influencing hyperpolarization. Interestingly, all 3 knockouts that resulted in severely hampered hyperpolarization are tied to redox stress and NADH/NAD+ ratio. Ndh directly controls redox by converting NADH into NAD+, while Rex represses genes when NADH/NAD+ is low (Härtig & Jahn, 2012) Together, Rex and Ndh provide a regulatory loop by which to control NADH/NAD+ ratio (Gyan et al., 2006). Meanwhile, in *B. amyloliquefaciens*, ResDE has been shown to be activated by an increase in NADH/NAD+ ratio (Zhou et al., 2018). The stark decrease in the ability of these mutants to hyperpolarize in response to oxygen downshift suggests a link between redox state and hyperpolarization. NADH/NAD+ ratio could play a role in inducing ΔΨ shifts during changing oxygen conditions, as has been previously suggested (Jones & Larkin, 2021). The fact that change in oxygen induces shifts in redox ratio (Härtig & Jahn, 2012) and ATP concentration (Supplementary Fig. 6C) in *B. subtilis,* coupled with the response of potassium transporters to these molecules (discussed above), provides a possible mechanism by which cells could hyperpolarize by changing ion flux during oxygen shift. A similar mechanism has been proposed for modulating ΔΨ in carotid bodies in response to acute hypoxia, where reactive oxygen species and NADH from mitochondria modulate potassium channels (Gao et al., 2017).

Hyperpolarization appears to play an important role in promoting cell proliferation following oxygen downshift, as evidenced by our finding that attenuation of hyperpolarization using valinomycin ionophore results in slower growth recovery. We theorize that cells release potassium during oxygen downshift as a mechanism for maintaining ΔΨ and PMF in the absence of proton pumping by the ETC. Krasnopeeva et al. recently highlighted the importance of both ATP and PMF in maintaining growth in *E. coli* (Krasnopeeva et al., 2026). While both fermentation and oxidative phosphorylation generate ATP, only oxidative phosphorylation contributes to PMF. Therefore, in anaerobic conditions where the cell relies on only fermentation, the ATP synthase F_1_F_O_ becomes essential for its reverse role in consuming ATP to pump out protons and establish PMF. We propose a similar mechanism where, in the absence of oxygen, the cell releases potassium to generate a strong enough ΔΨ to remain viable and carry out crucial processes that rely on ΔΨ, such as nutrient acquisition, particularly glutamate. In *B. subtilis*, glutamate can be imported through the GltP symporter in a ΔΨ-dependent manner (Tolner et al., 1995). Glutamate has been suggested to play a role in oxidative stress tolerance through the production of the antioxidant glutathione (Addorisio et al., 2022; Feehily & Karatzas, 2013). For example, glutathione production has been shown to increase the survival of *Salmonella* in macrophages (Huang et al., 2022). We propose that membrane hyperpolarization during oxygen downshift aids in glutamate uptake to provide protection from oxidative stress. Hyperpolarizing in response to oxygen downshift may enhance fitness in dynamic environments by ensuring that crucial ΔΨ-reliant processes keep running so that cells remain viable and growth may rapidly resume once oxygen levels are restored.

Oxygen-induced hyperpolarization is also of clinical relevance, as the uptake of antibiotics, specifically aminoglycosides, relies on ΔΨ (Bruni & Kralj, 2020; Damper & Epstein, 1981; Ezraty et al., 2013). Our finding that oxygen shift increased bactericidal activity of antibiotics presents the potential for coordinating drug application with oxygen conditions as an approach to improve treatment efficacy.

Membrane potential and aerobic respiration sit at the nexus of numerous physiological processes, including metabolism, redox state, osmoregulation and stress response. Cellular hyperpolarization in response to oxygen drop, therefore, has broad significance for cell physiology. Our findings open questions about how oxygen shifts are sensed and transduced into changes in ion transporter activity, and whether the response could subsequently result in broader changes to cell programming, such as altered gene expression, to aid in long term survival in hypoxia. Furthermore, the generation of membrane potential using oxygen as an electron acceptor has defined the evolution of aerobic life. The conserved hyperpolarization response we find in aerobic bacteria parallels mitochondrial dynamics during hypoxia, raising questions on the evolution of microbial electrophysiology and its impact on the development of multicellularity.

## Terms

ΔΨ: membrane potential
PMF: proton motive force
ETC: electron transport chain
ΔThT: net change in ThT signal after 1 hour of oxygen drop
fFOV: fraction of the field of view covered by cell monolayer
ΔThTp: predicted ΔThT at .6 fFOV
Vmax: maximum velocity of fFOV expansion

## Supplementary Figure captions

**Supplementary Fig. 1.**
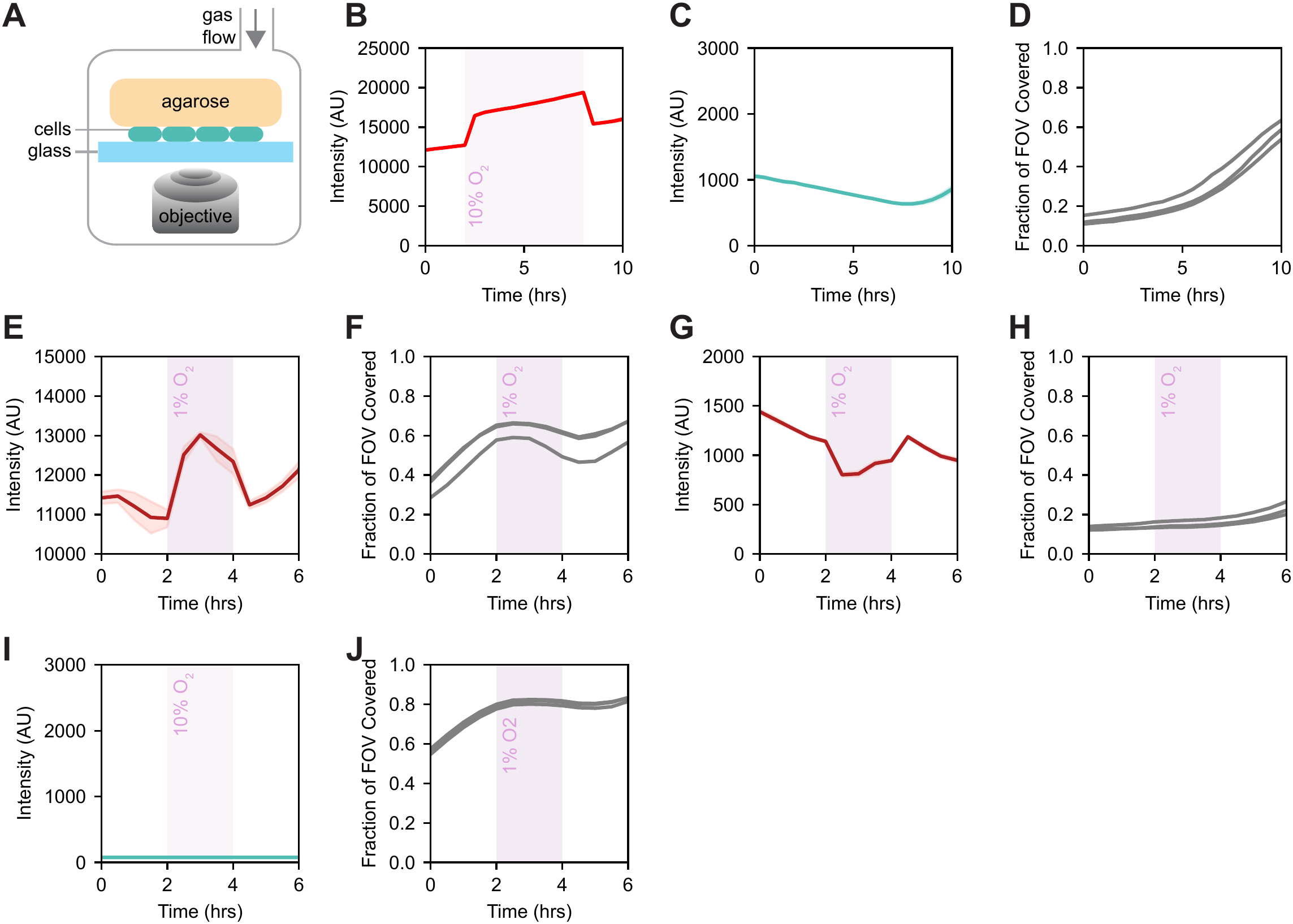
Bacteria hyperpolarize during oxygen downshift and depolarize upon oxygen upshift. **A.** Schematic for agarose pad imaging and gas control setup. Cells are sandwiched between agarose and glass and imaged from below. Gas with desired concentration of oxygen is flowed into the enclosed chamber. **B.** RTDP oxygen reporter in the absence of cells showing the shift in oxygen in the agarose when oxygen shifts in the chamber. **C.** *B. subtilis* Mean ThT fluorescence intensity over 10 hours with no oxygen shift for 3 fields of view. Shaded region indicates mean +/- s. e. m. **D.** fFOV over time for 3 fields of view measured in (C). **E**. *B. subtilis* mean TMRM fluorescence intensity over time for 3 fields of view. Shaded region around the line indicates mean +/- s. e. m. **F.** fFOV over time for 3 fields of view measured in (E). **G.** *B. subtilis* mean DISC(3)5 fluorescence intensity over time for 3 fields of view. Note that DISC3(5) behaves inversely to ThT, with a decrease in fluorescence indicating hyperpolarization. Shaded region indicates mean +/- s. e. m. **H.** fFOV over time for 3 fields of view measured in (G). **I.** Mean ThT fluorescence intensity in the absence of cells over time for 3 fields of view. Shaded region indicates s. e. m. **J.** *B. subtilis* fFOV over time for 3 fields of view in the absence of ThT for 3 fields of view. Shaded region indicates mean +/- s. e. m.

**Supplementary Fig. 2.**
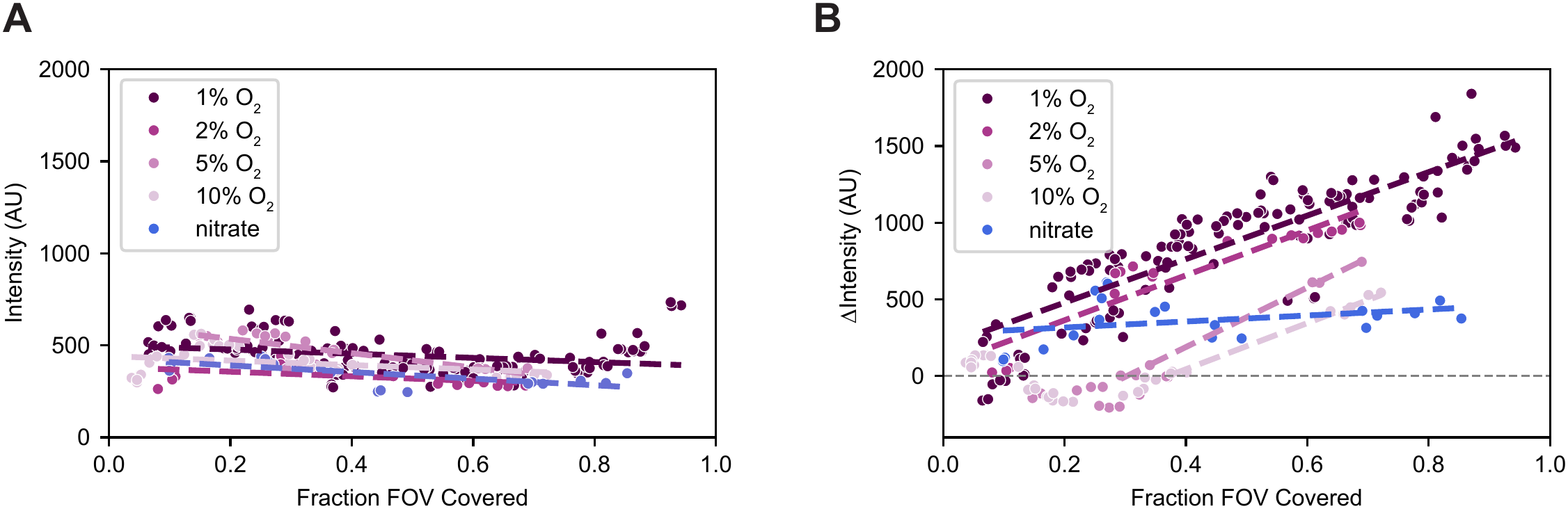
Hyperpolarization magnitude scales with the degree oxygen deprivation. **A.** Scatterplot for raw ThT intensity before oxygen downshift to either 1%, 2%, 5% or 10% oxygen versus fFOV. Nitrate indicates cells that undergo a 1% oxygen downshift on media supplemented with nitrate. Each point represents one field of view imaged. Dashed lines represent linear regression. **B.** Scatterplot for change in ThT intensity after 1hr of oxygen downshift to either 1%, 2%, 5% or 10% oxygen versus fFOV at the time of oxygen downshift, as seen in Fig. 2E but including the nitrate condition. Each point represents one field of view imaged. Dashed lines represent linear regression for points with positive net intensity and to the right of the first negative Δintensity value.

**Supplementary Fig. 3.**
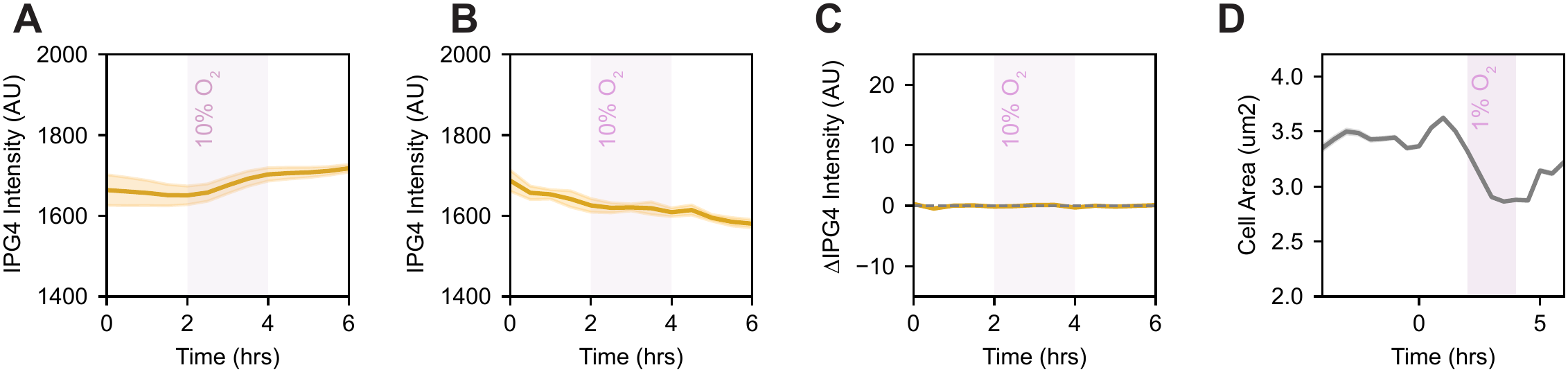
Potassium efflux contributes to hyperpolarization. **A.** Mean IPG-4 extracellular potassium dye fluorescence intensity over time for 3 different fields of view. Shaded region indicates mean +/- s. e. m. Pink block indicates the duration of 10% oxygen. **B.** Mean IPG-4 extracellular potassium dye fluorescence intensity over time in the absence of cells for 3 different fields of view. Shaded region indicates mean +/- s. e. m. Pink block indicates the duration of 10% oxygen. **C.** Mean change in IPG-4 fluorescence intensity over time from (B). Shaded region indicates mean +/- s. e. m. Pink block indicates the duration of 10% oxygen. **D.** Mean single cell area over time in the absence of ThT for 3 different fields of view. Shaded region indicates mean +/- s. e. m. Pink block indicates the duration of 1% oxygen.

**Supplementary Fig. 4.**
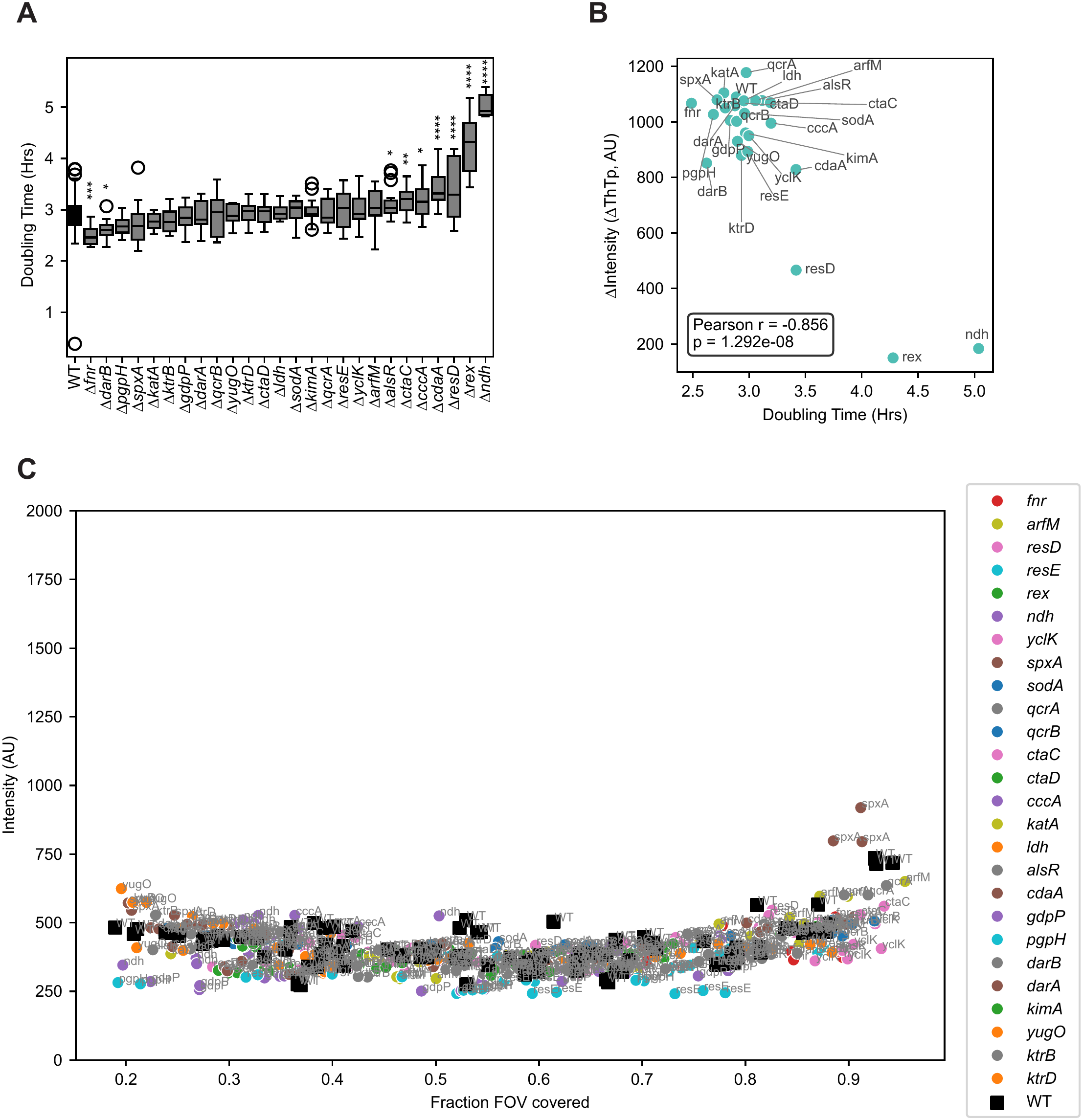
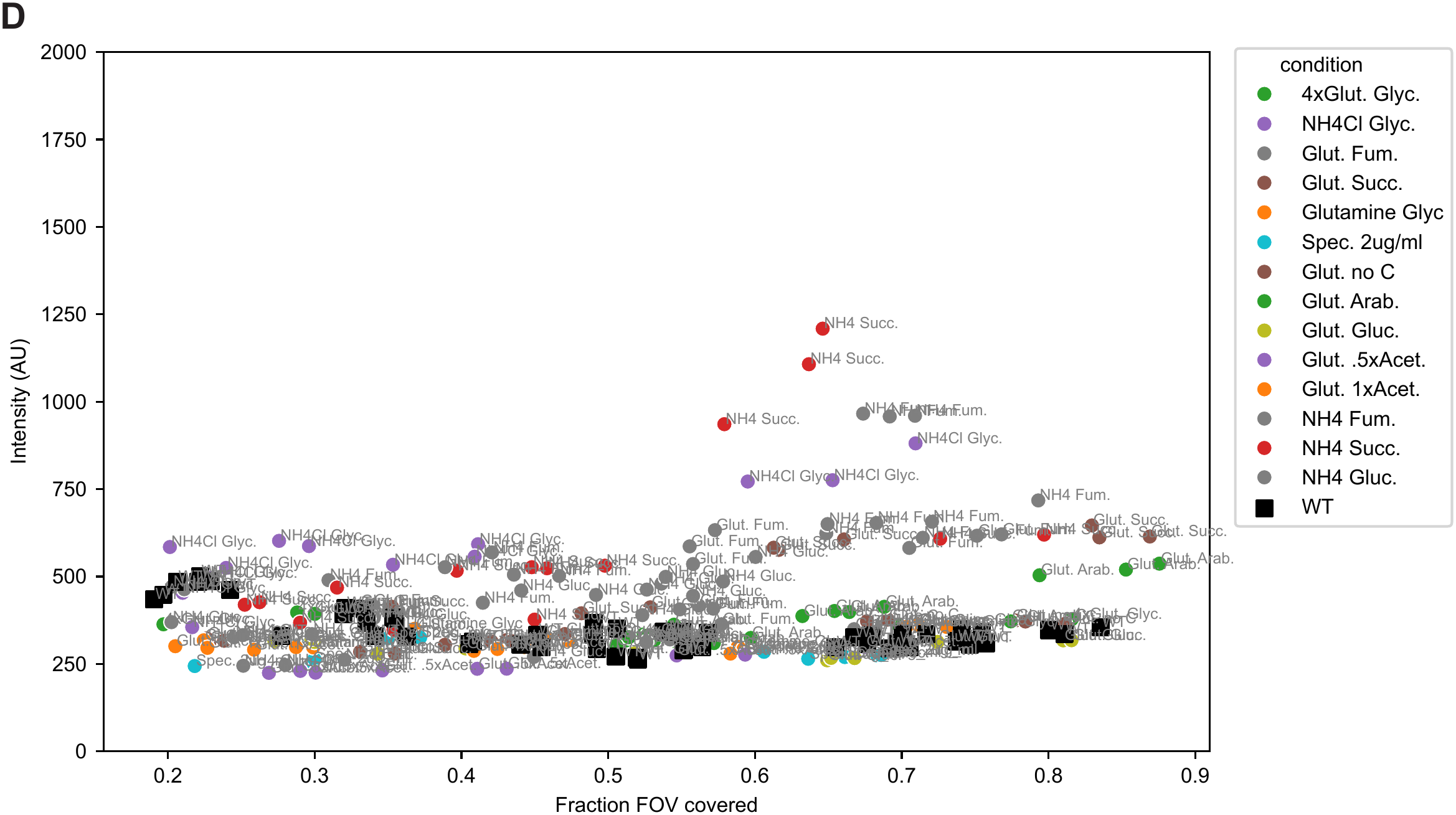
Oxygen stress genes impact hyperpolarization response to oxygen downshift. **A.** Mean doubling time in 21% oxygen (air) for different gene knockouts of *B. subtilis*. Error bars represent standard error. N ≥ 4 for each condition. **B.** Scatterplot for predicted change in ThT intensity after 1 hour of oxygen downshift to 1% oxygen at .6 fFOV (shown in Fig. 4B) versus the average doubling time for each nutrient/antibiotic in 21% oxygen (air) (shown here in A). Pearson r shows the Pearson correlation and p shows the p-value. **C.** Scatterplot for raw ThT intensity before oxygen downshift versus fFOV for different gene knockouts. Each point represents one field of view imaged. **D.** Scatterplot for raw ThT intensity before oxygen downshift versus fFOV for different media compositions. Each point represents one field of view imaged.

**Supplementary Fig. 5.**
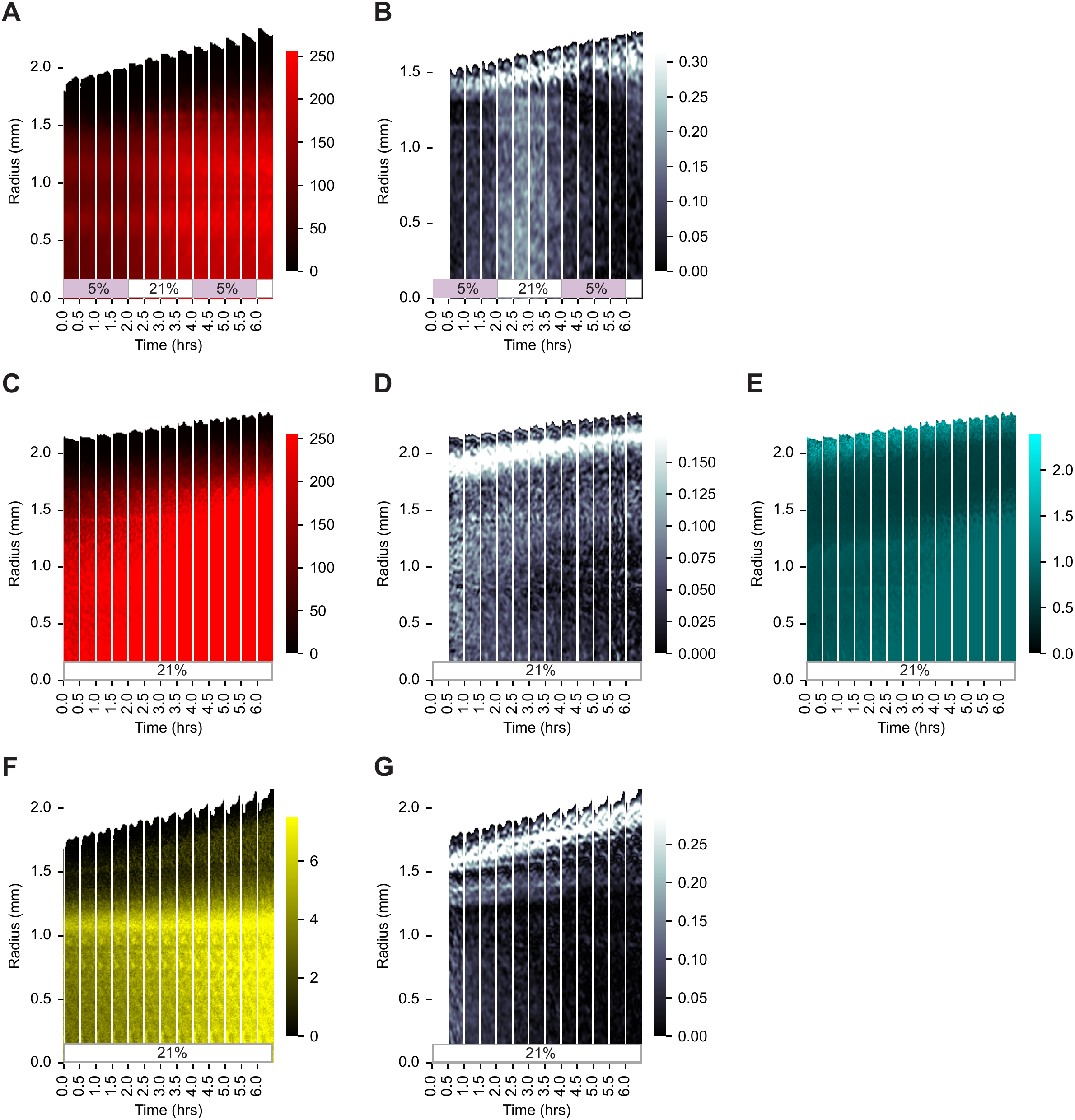
Magnitude of hyperpolarization scales with growth rate in both microcolonies and biofilms. **A, B.** Radial strips of 18 hr-old biofilms undergoing oxygen shifts from 5% oxygen to 21% (air) every 2 hours. **A.** Constitutive mScarlet signal for biofilm shown in Fig. 5D. **B.** phase difference for biofilm shown in Fig. 5F. **C-E.** Radial strips of an 18 hr-old biofilm not undergoing oxygen shifts shown in constitutive mScarlet (C), phase difference (D) and normalized ThT (E). **F**, **G.** Radial strips of an 18 hr-old biofilm not undergoing oxygen shifts shown in IPG-4 extracellular potassium dye (**F**) and phase difference (**G**).

**Supplementary Fig. 6.**
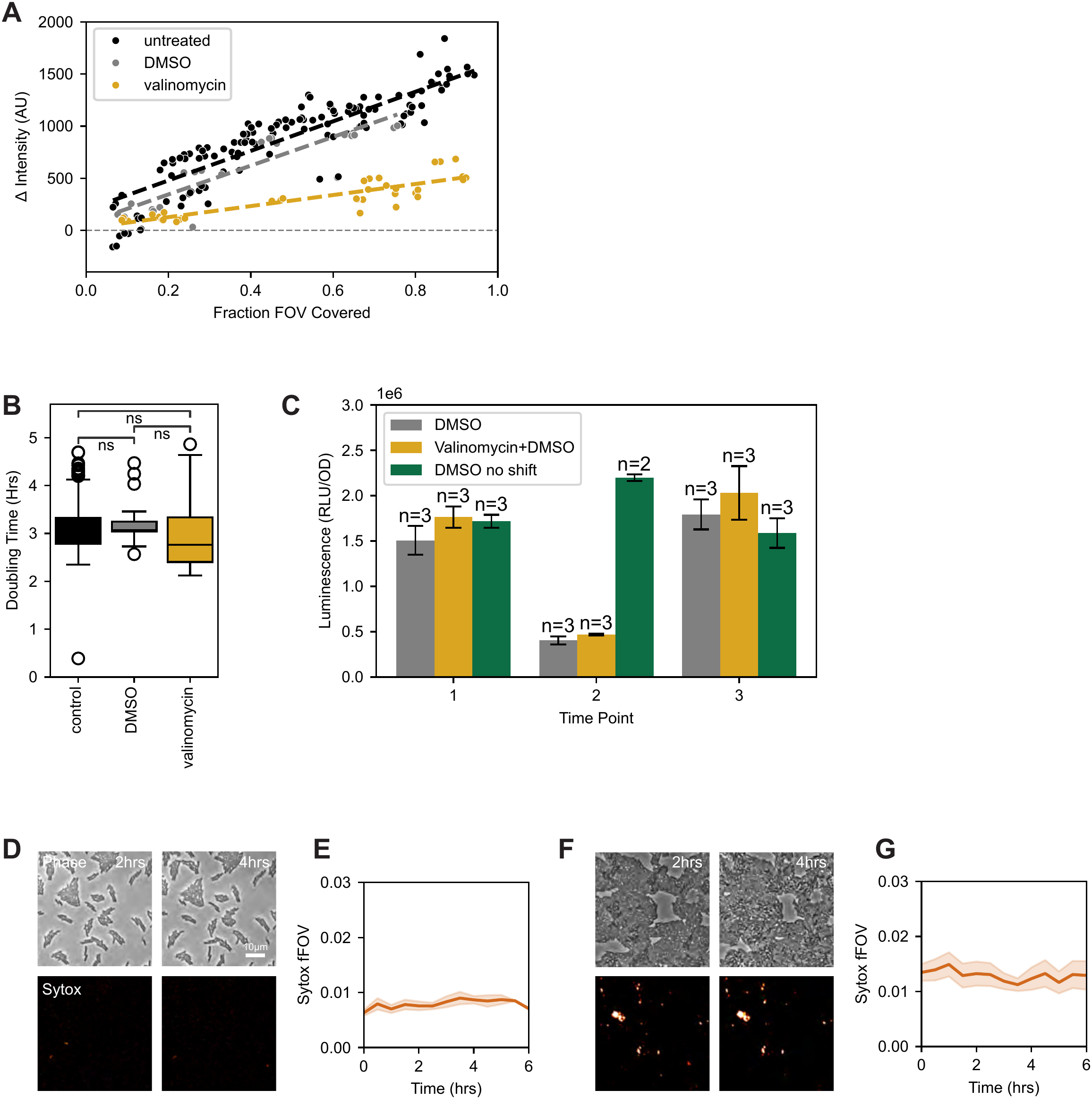
Hyperpolarization supports cell proliferation and increases antibiotic susceptibility. **A.** Scatterplot for change in ThT intensity after 1hr of oxygen downshift to either untreated, DMSO control or valinomycin, versus fFOV at the time of oxygen downshift. Each point represents one field of view imaged. Dashed lines represent linear regression. **B.** Mean doubling time in either DMSO control or valinomycin for WT *B. subtilis* in the absence of oxygen shifts. Error bars represent standard error. N ≥ 6 for each condition. Stars indicate significance (* p value <.05, ** p value <.01, *** p value <.001). **C.** ATP level measured by luciferase assay for *B. subtilis* in either untreated, DMSO control or valinomycin, before oxygen shift (time point 1), after 1hr in anaerobic conditions (time point 2), and after a 1hr recovery period in air (time point 3). Green represents control condition of DMSO that did not undergo oxygen shift. *B. subtilis* with no oxygen shift in air (21% oxygen) with kanamycin (**D-E**), or spectinomycin (**F-G**). D, F. still images of cells at 2 hrs (left) and 4 hrs (right) in phase (top) and Sytox Orange cell death stain (bottom). E, G. Mean fraction of field of view covered by Sytox signal (Sytox fFOV) over time. N = 3 fields of view. Shaded region indicates mean +/- s. e. m.

## Movies

**Movie 1:** Fluorescence signal of the membrane potential reporter ThT in a *B. subtilis* microcolony on an agarose pad undergoing an oxygen shift from air (21%) to 1% between 2-4 hrs. Time stamp is in hh:mm.

**Movie 2:** Phase imaging of a *B. subtilis* microcolony on an agarose pad undergoing an oxygen shift from air (21%) to 1% between 2-4 hrs. Time stamp is in hh:mm.

**Movie 3:** Fluorescence signal of the membrane potential reporter ThT normalized to constitutive mScarlet reporter of an 18 hr-old *B. subtilis* biofilm undergoing oxygen shifts from 5% oxygen to 21% (air) every 2 hours. Time stamp is in hh:mm.

**Movie 4:**

Phase difference calculation of an 18 hr-old *B. subtilis* biofilm (shown in Movie 3) undergoing oxygen shifts from 5% oxygen to 21% (air) every 2 hours. Time stamp is in hh:mm.

## Tables

**Supplementary Table 1.**

| strain<br>identifier | species | mutation | description |
|---|---|---|---|
| MK0 | *B. subtilis* |  | 3610 (WT) |
| JWL6 | *P. megaterium* |  |  |
| MK23 | *E. coli* |  | MG1655 |
| MK25 | *M. luteus* |  |  |
| MK32 | *B. subtilis* | Δ*darB* | darB::erm |
| MK33 | *B. subtilis* | Δ*darA* | darA::erm |
| MK34 | *B. subtilis* | Δ*kimA* | kimA::erm |
| MK4 | *B. subtilis* | Δ*cdaA* | cdaA::erm |
| MK8 | *B. subtilis* | Δ*gdpP* | gdpP::erm |
| MK10 | *B. subtilis* | Δ*pgpH* | pgpH::erm |
| MK147 | *B. subtilis* | Δ*fnr* | fnr::erm |
| MK148 | *B. subtilis* | Δ*arfM* | arfM::erm |
| MK149 | *B. subtilis* | Δ*resD* | resD::erm |
| MK150 | *B. subtilis* | Δ*resE* | resE::erm |
| MK151 | *B. subtilis* | Δ*rex* | rex::erm |
| MK152 | *B. subtilis* | Δ*ndh* | ndh::erm |
| MK153 | *B. subtilis* | Δ*yclK* | yclK::erm |
| MK154 | *B. subtilis* | Δ*ldh* | ldh::erm |
| MK155 | *B. subtilis* | Δ*alsR* | alsR::erm |
| MK160 | *B. subtilis* | Δ*yugO* | yugO::erm |
| MK161 | *B. subtilis* | Δ*ktrB* | ktrB::erm |
| MK162 | *B. subtilis* | Δ*ktrD* | ktrD::erm |
| JMJ1217 | *B. subtilis* | Constitutive mScartlet | amyE::Pveg-mScarlet-spec |

**Supplementary Table 2.**
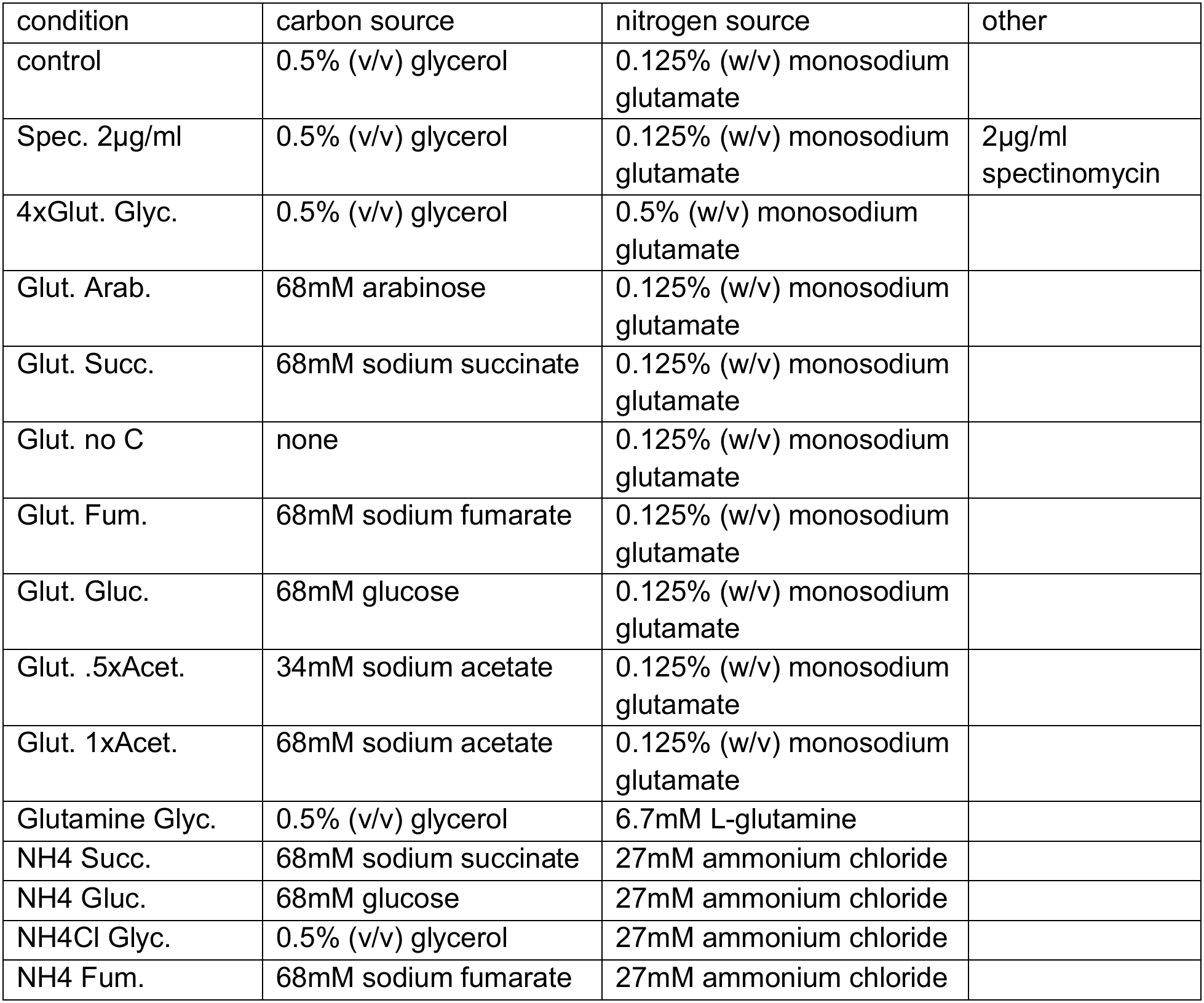

| condition | carbon source | nitrogen source | other |
|---|---|---|---|
| control | 0.5% (v/v) glycerol | 0.125% (w/v) monosodium glutamate |  |
| Spec. 2µg/ml | 0.5% (v/v) glycerol | 0.125% (w/v) monosodium glutamate | 2µg/ml spectinomycin |
| 4xGlut. Glyc. | 0.5% (v/v) glycerol | 0.5% (w/v) monosodium glutamate |  |
| Glut. Arab. | 68mM arabinose | 0.125% (w/v) monosodium glutamate |  |
| Glut. Succ. | 68mM sodium succinate | 0.125% (w/v) monosodium glutamate |  |
| Glut. no C | none | 0.125% (w/v) monosodium glutamate |  |
| Glut. Fum. | 68mM sodium fumarate | 0.125% (w/v) monosodium glutamate |  |
| Glut. Gluc. | 68mM glucose | 0.125% (w/v) monosodium glutamate |  |
| Glut. .5xAcet. | 34mM sodium acetate | 0.125% (w/v) monosodium glutamate |  |
| Glut. 1xAcet. | 68mM sodium acetate | 0.125% (w/v) monosodium glutamate |  |
| Glutamine Glyc. | 0.5% (v/v) glycerol | 6.7mM L-glutamine |  |
| NH4 Succ. | 68mM sodium succinate | 27mM ammonium chloride |  |
| NH4 Gluc. | 68mM glucose | 27mM ammonium chloride |  |
| NH4Cl Glyc. | 0.5% (v/v) glycerol | 27mM ammonium chloride |  |
| NH4 Fum. | 68mM sodium fumarate | 27mM ammonium chloride |  |

## Methods

### Strains

The *B. subtilis* strains used are listed in Supplementary Table 1. Standard techniques were used for cloning and strain construction (Harwood & Cutting, 1990). We used a transformable derivative of strain NCIB3610 (Konkol et al., 2013), which was obtained from the Bacillus Genetic Stock Center (BGSC).

### Construction of knockout mutants

Gene knockouts were generated by allelic replacement of the indicated genes with antibiotic resistance genes. Homology arms (∼2 kb) upstream and downstream of the indicated open reading frames (ORF) were cloned and fused to an antibiotic resistance gene cassette by isothermal assembly (Gibson et al., 2009). The resulting DNA fragment was introduced to *B. subtilis* by transformation and antibiotic selection.

### Construction of constitutive reporter

To label cellular biomass, a strong transcriptional unit composed of the Pveg promoter and R0 ribosome binding site (Guiziou et al., 2016) was used to drive expression of mScarlet (Bindels et al., 2017). The Pveg-R0-mScarlet fragment was fused to a spectinomycin resistance cassette and homology arms for integration at *amyE* as previously described.

### Media

Prior to experiments, cells were grown as light lawns for approximately 24 hours at room temperature on 1.5% agar plates containing 1% w/v glucose, 0.1% w/v monosodium glutamate, and 1x Spizizen’s salts (2 g/l (NH_4_)SO_4_, 14 g/l K_2_HPO_4_, 6 g/l KH_2_PO_4_, 1 g/l Na_3_citrate-2H_2_O, and 0.2 g/l MgSO_4_-7H_2_O) (Harwood & Cutting, 1990). Light lawns were used to bypass liquid culture steps commonly used to prepare cells, as this selects for biofilm-deficient mutants.

Experiments were done on MSgg media composed of 5 mM potassium phosphate (pH 7.0), 100 mM MOPS (pH 7.0), 2 mM MgCl2, 700 μM CaCl2, 50 μM MnCl2, 100 μM FeCl3, 1 μM ZnCl2, 2 μM thiamine HCl, 0.5% (v/v) glycerol and 0.125% (w/v) monosodium glutamate. MSgg media was prepared on the day of the experiment from stock solutions, with glutamate and iron stock solutions prepared the day of. Variations to media composition for Figure 5 can be seen in Supplementary Table 2.

Agarose pads were prepared with the designated media and 3% agarose and relevant fluorescent dyes. 916 µl mixture was sandwiched between two 22 mm x 22 mm cover slips and allowed to dry for 2 hours. These were then quartered using a razor blade and 5 µl of cells (diluted from light lawns) were spotted into the center of each. Pads were then flipped upside down onto 24-well plates (Cellvis) with one pad per well. 24-well plates were sealed with Breathe-Easy film to prevent evaporation and 2 holes were poked into the film at each well to allow for gas flow. Plates were allowed to acclimate for 6 hrs in air (∼21% oxygen) and then subjected to a 2 hr oxygen drop before returning to air once again. The experiments shown begin 4 hours into the acclimation period, such that they begin 2 hours before the oxygen drop.

Colony experiments were performed in 24-well plates, each well with 400 µl of 3% agarose and MSgg mixture. 1µl of cells at OD .5 (diluted from light lawns) was spotted into wells and plates were then sealed with Breathe-Easy film to prevent evaporation. 2 holes were poked into the film at each well to better allow for gas flow. Plates were then subjected to 2-hour cycles of air and 5% oxygen. Colony data is shown in Figure 6D-F begins 18 hrs into the experiment, starting at 5% oxygen.

### Dyes, ionophores and antibiotics

Fluorescent dyes were used to visualize membrane potential. ThT and DISC(3)5 were used at 10 μM, and TMRM at .1 μM. IPG-4 TMA+ Salt from ION Biosciences was used at 2 μM for visualizing extracellular potassium. Ruthenium tris(2,2’-dipyridyl) dichloride hexahydrate (RTDP) was used at 0.25 mg/mL to visualize changes in oxygen. The ionophore valinomycin was used to disturb membrane potential and was used at 5 µM and the same concentration of DMSO was used as a control. For cell death experiments, Sytox Orange was used at 1 µM and the antibiotics spectinomycin and kanamycin were used at 3 mg/L and .07 mg/L respectively for cell death experiments.

### Imaging and gas control

Experiments were run at 30°C and oxygen content was regulated using a gas mixer (Okolab, Pozzuoli, Italy) to mix air and nitrogen gas. Gas cycles were automated using DATALOG. Agarose pads of cell monolayer and biofilm colonies were imaged with a 40X, 0.75 NA and 4X 0.13 NA objective respectively, in phase contrast and fluorescence on an Olympus IX-83 microscope (Evident Scientific, Waltham, MA, USA). At 40X, ThT was imaged at 25 ms using a CFP filter cube, DISC(3)5 and TMRM at 25 ms using an RFP filter cube, IPG-4 at 100 ms using a YFP filter cube, and Sytox Orange at 15 ms using an RFP filter cube. At 4X, ThT was imaged at 25 ms using a CFP filter cube, and IPG-4 at 100ms using a YFP filter cube.

### Image analysis, ΔThT and fFOV and ΔThTp

Time traces of fluorescent dyes were extracted from time-lapse movies using Weka Segmentation in ImageJ and custom python scripts using phase contrast images for segmentation. Signal was determined by taking the mean signal of the cell-covered region for 3 fields of view on the same agarose pad. fFOV was defined as the corresponding fraction of the field of view covered by cell monolayer. Change in intensity (ΔThT) was calculated by subtracting the ThT intensity at 2 hrs (immediately before oxygen drop) from the ThT intensity at 3 hrs (1 hr into the oxygen drop). Predicted change in ThT intensity (ΔThTp) was calculated by taking the linear regression of the scatter plot of ΔThT versus fFOV for a given condition and reporting its value and 95^th^% confidence interval at .6 fFOV. Linear regression was derived using bootstrap resampling. Mean parameter estimates and two-sided 95% confidence intervals were calculated from the bootstrap distributions.

### Doubling time calculation

Doubling times were estimated from the measured fFOV trajectories in 21% oxygen (air). The initial fFOV was used to define a twofold-increased target fFOV. The time at which this target was reached was estimated by log-linear interpolation between the two consecutive measurements bracketing the target, assuming exponential growth between measurements. Doubling time was calculated as the elapsed time between the initial measurement and the interpolated time at which the fFOV reached twice its initial value.

### Cell size measurements

Cell area was calculated by performing single-cell segmentation on phase images using Omnipose (Cutler et al., 2022) and taking the mean cell size using custom python code.

### Vmax

Vmax was calculated as the vertex of the quadratic regression of the change in fFOV (ΔfFOV/Δt) for different fFOVs at a given time point.

### Colony analysis

Biofilm colonies were segmented using custom python code. ThT signal was normalized by dividing ThT intensity by constitutively expressed mScarlet intensity. Phase difference was calculated by subtracting phase images by the previous image. For IPG-4 and mScarlet, raw intensity values are shown.

### ATP Measurements

ATP was measured using Promega BacTiter-Glo kit and following the kit protocol. Cells were evenly plated on MSgg and agar 12-well plates with either valinomycin or DMSO control and grown at 30°C. Plates were sealed with Breathe-Easy film to prevent evaporation and 2 holes were poked into the film at each well to allow for gas flow. At 22 hrs, cells were subjected to low oxygen by being placed in an anaerobic chamber for 1hr and then allowed to recover in air for 1 hr. Cells were collected by washing off with MSgg media before and at the end of oxygen shift as well as after recovering 1 hr in air. The low oxygen sample was collected with MSgg that had been sparged for 20 min with nitrogen gas. Bioluminescence and OD was measured in 96-well plates using a Synergy H1 microplate reader (Agilent Biotek) and bioluminescence signal was normalized by dividing by the OD of cells.

### Steady-state reaction-diffusion model

The method of computationally evaluating steady-state oxygen competition contains 4 steps: establishing colony geometry, defining physical conditions, solving uptake rates, and mapping uptake onto a hyperpolarization response. Our model assumes that after an external shift, oxygen reaches a new steady-state on a faster time scale than the hour-long measurement; oxygen diffuses across a 75 µm range of cell–cell interactions within seconds. The model also treats the colony geometry as fixed during the measurement, which is supported by the observation that growth stalls in low oxygen (Fig. 1C).

Establishing colony geometry uses a modified hard-core Poisson disk process. Cells are represented as disks of radius a =1 μm, and cell centers are randomly generated with a minimum center-to-center separation of 2a to prevent cell overlap. The probability that a newly initialized cell will join an existing cluster is set to 60%. This produces clusters that qualitatively resemble the spatial organization observed in experimental images.

Because the model describes the steady-state oxygen field, oxygen concentration in the extracellular region is governed by the two-dimensional steady-state diffusion equation. The physical condition at each cell is a type of Robin boundary condition (Collins & Kimball, 1949). It specifies that the uptake flux density *J_i_* is proportional to the local concentration at the cell boundary, with coefficient *κ*:

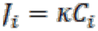

The flux density for cell *i* can be multiplied by the cell’s 2D circumference (2π*a*) to obtain the cell-specific uptake quantity *q_i_*.

To solve for the cell-specific uptake rates, cell–cell interactions were represented using a truncated Green’s function derived from the 2D steady state diffusion equation (Laplace equation), where *r* is the distance between two cell centers:

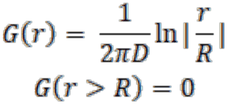

The interaction kernel is truncated to 0 beyond a cutoff distance *R*, omitting the contributions of more distant cells. A center-to-center cutoff of *R* = 75 µm was used for the reported simulations.

Next, cell-specific uptake quantities are calculated by solving a linear system defined below, where N is the total number of cells and *C_∞_* represents the background dissolved oxygen concentration in the pad, in equilibrium with the imposed gas-phase oxygen percentage.

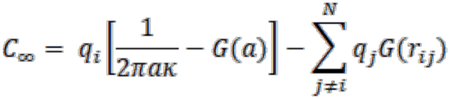

Finally, cell-specific oxygen uptake was mapped onto a normalized hyperpolarization response. Cells with uptake above a critical threshold were assigned no response; below this threshold, the response increased linearly as uptake decreased, reaching a value of one at zero uptake. The population-level simulated response was calculated as the mean hyperpolarization response across cells, consistent with the experimental measurement of the mean change in fluorescence intensity over cell-covered regions.

Model parameters were fitted by minimizing the mean, over the four oxygen levels, of the root-mean-square error between simulated and measured responses (SI §S6). No single fixed threshold reproduced all four oxygen levels. The threshold was therefore allowed to increase linearly with the oxygen level after the shift, f_c_ = f₀ + α O_env_, where O_env_ is the oxygen level in percent.

## Acknowledgements

Wild-type *B. subtilis* NCIB3610 and *P. megaterium* were obtained from the Bacillus Genetic Stock Center. *M. luteus* and *E. coli* were gifts from Zeba Wunderlich and Mary Dunlop, respectively. The constitutive *mScarlet* reporter was constructed by Joshua Jones. We acknowledge members of the Larkin Lab for useful discussions and Joshua Jones, Emily Hager, Chiara Ricci-Tam, Xinwen Zhu, and Caroline Ajo-Franklin for all their helpful insights.

## References

Addorisio, S. R., Shteynberg, R. M., Dasilva, M. S., Mixon, J. M., Mucciarone, K., Vu, L., Arsenault, K. L., Briand, V., Parker, S., Smith, S. L., Vise, C. E., Pina, C., & Laranjo, L. T. (2022). Oxidative Stress Response in Bacteria: A Review.

Arjes, H. A., Vo, L., Dunn, C. M., Willis, L., DeRosa, C. A., Fraser, C. L., Kearns, D. B., & Huang, K. C. (2020). Biosurfactant-Mediated Membrane Depolarization Maintains Viability during Oxygen Depletion in Bacillus subtilis. Current Biology, 30(6), 1011–1022.e6. 10.1016/j.cub.2020.01.073

Berry, B. J., Trewin, A. J., Amitrano, A. M., Kim, M., & Wojtovich, A. P. (2018). Use the Protonmotive Force: Mitochondrial Uncoupling and Reactive Oxygen Species. In Journal of Molecular Biology (Vol. 430, Number 21, pp. 3873–3891). Academic Press. 10.1016/j.jmb.2018.03.025

Bindels, D. S., Haarbosch, L., van Weeren, L., Postma, M., Wiese, K. E., Mastop, M., Aumonier, S., Gotthard, G., Royant, A., Hink, M. A., & Gadella, T. W. J. (2017). mScarlet: a bright monomeric red fluorescent protein for cellular imaging. Nature Methods, 14(1), 53–56. 10.1038/nmeth.4074

Biquet-Bisquert, A., Labesse, G., Pedaci, F., & Nord, A. L. (2021). The Dynamic Ion Motive Force Powering the Bacterial Flagellar Motor. Frontiers in Microbiology, 12. 10.3389/fmicb.2021.659464

Brogan, A. P., Bardetti, P., Rojas, E. R., & Rudner, D. Z. (2025). Cyclic-di-AMP modulates cellular turgor in response to defects in bacterial cell wall synthesis. Nature Microbiology, 10(7), 1698–1710. 10.1038/s41564-025-02027-2

Bruni, G. N., & Kralj, J. M. (2020). Membrane voltage dysregulation driven by metabolic dysfunction underlies bactericidal activity of aminoglycosides. eLife, 9. 10.7554/eLife.58706

Bruni, G. N., Weekley, R. A., Dodd, B. J. T., & Kralj, J. M. (2017). Voltage-gated calcium flux mediates Escherichia coli mechanosensation. Proceedings of the National Academy of Sciences of the United States of America, 114(35), 9445–9450. 10.1073/pnas.1703084114

Bueno, E., Mesa, S., Bedmar, E. J., Richardson, D. J., & Delgado, M. J. (2012). Bacterial adaptation of respiration from oxic to microoxic and anoxic conditions: Redox control. In Antioxidants and Redox Signaling (Vol. 16, Number 8, pp. 819–852). 10.1089/ars.2011.4051

Cao, Y., Pan, Y., Huang, H., Jin, X., Levin, E. J., Kloss, B., & Zhou, M. (2013). Gating of the TrkH ion channel by its associated RCK protein TrkA. Nature, 496(7445), 317–322. 10.1038/nature12056

Choi, H., Yang, Z., & Weisshaar, J. C. (2017). Oxidative stress induced in E. coli by the human antimicrobial peptide LL-37. PLOS Pathogens, 13(6), e1006481. 10.1371/journal.ppat.1006481

Collins, F. C., & Kimball, G. E. (1949). Diffusion-controlled reaction rates. Journal of Colloid Science, 4(4), 425–437. 10.1016/0095-8522(49)90023-9

Commichau, F. M., Gibhardt, J., Halbedel, S., Gundlach, J., & Stülke, J. (2018). A Delicate Connection: c-di-AMP Affects Cell Integrity by Controlling Osmolyte Transport. In Trends in Microbiology (Vol. 26, Number 3, pp. 175–185). Elsevier Ltd. 10.1016/j.tim.2017.09.003

Cutler, K. J., Stringer, C., Lo, T. W., Rappez, L., Stroustrup, N., Brook Peterson, S., Wiggins, P. A., & Mougous, J. D. (2022). Omnipose: a high-precision morphology-independent solution for bacterial cell segmentation. Nature Methods, 19(11), 1438–1448. 10.1038/s41592-022-01639-4

Damper, P. D., & Epstein, W. (1981). Role of the membrane potential in bacterial resistance to aminoglycoside antibiotics. Antimicrobial Agents and Chemotherapy, 20(6), 803–808. 10.1128/AAC.20.6.803

Ezraty, B., Vergnes, A., Banzhaf, M., Duverger, Y., Huguenot, A., Brochado, A. R., Su, S.-Y., Espinosa, L., Loiseau, L., Py, B., Typas, A., & Barras, F. (2013). Fe-S Cluster Biosynthesis Controls Uptake of Aminoglycosides in a ROS-Less Death Pathway. Science, 340(6140), 1583–1587. 10.1126/science.1238328

Feehily, C., & Karatzas, K. A. G. (2013). Role of glutamate metabolism in bacterial responses towards acid and other stresses. Journal of Applied Microbiology, 114(1), 11–24. 10.1111/j.1365-2672.2012.05434.x

Gao, L., González-Rodríguez, P., Ortega-Sáenz, P., & López-Barneo, J. (2017). Redox signaling in acute oxygen sensing. Redox Biology, 12, 908–915. 10.1016/j.redox.2017.04.033

Geng, H., Nakano, S., & Nakano, M. M. (2004). Transcriptional Activation by Bacillus subtilis ResD: Tandem Binding to Target Elements and Phosphorylation-Dependent and - Independent Transcriptional Activation. Journal of Bacteriology, 186(7), 2028–2037. 10.1128/JB.186.7.2028-2037.2004

Gibson, D. G., Young, L., Chuang, R.-Y., Venter, J. C., Hutchison, C. A., & Smith, H. O. (2009). Enzymatic assembly of DNA molecules up to several hundred kilobases. Nature Methods, 6(5), 343–345. 10.1038/nmeth.1318

Gray, D. A., Wang, B., Sidarta, M., Cornejo, F. A., Wijnheijmer, J., Rani, R., Gamba, P., Turgay, K., Wenzel, M., Strahl, H., & Hamoen, L. W. (2024). Membrane depolarization kills dormant Bacillus subtilis cells by generating a lethal dose of ROS. Nature Communications, 15(1). 10.1038/s41467-024-51347-0

Guiziou, S., Sauveplane, V., Chang, H.-J., Clerté, C., Declerck, N., Jules, M., & Bonnet, J. (2016). A part toolbox to tune genetic expression in *Bacillus subtilis*. Nucleic Acids Research, gkw624. 10.1093/nar/gkw624

Gundlach, J., Herzberg, C., Hertel, D., Thürmer, A., Daniel, R., Link, H., & Stülke, J. (2017). Adaptation of *Bacillus subtilis* to Life at Extreme Potassium Limitation. mBio, 8(4). 10.1128/mBio.00861-17

Gundlach, J., Herzberg, C., Kaever, V., Gunka, K., Hoffmann, T., Weiß, M., Gibhardt, J., Thürmer, A., Hertel, D., Daniel, R., Bremer, E., Commichau, F. M., & Stülke, J. (2017). Control of potassium homeostasis is an essential function of the second messenger cyclic di-AMP in *Bacillus subtilis*. Science Signaling, 10(475). 10.1126/scisignal.aal3011

Gyan, S., Shiohira, Y., Sato, I., Takeuchi, M., & Sato, T. (2006). Regulatory loop between redox sensing of the NADH/NAD+ ratio by Rex (YdiH) and oxidation of NADH by NADH dehydrogenase Ndh in Bacillus subtilis. Journal of Bacteriology, 188(20), 7062–7071. 10.1128/JB.00601-06

Harold, F. M. (1972). Conservation and transformation of energy by bacterial membranes. Bacteriological Reviews, 36(2), 172–230. 10.1128/br.36.2.172-230.1972

Härtig, E., Geng, H., Hartmann, A., Hubacek, A., Münch, R., Ye, R. W., Jahn, D., & Nakano, M. M. (2004). *Bacillus subtilis* ResD Induces Expression of the Potential Regulatory Genes *yclJK* upon Oxygen Limitation. Journal of Bacteriology, 186(19), 6477–6484. 10.1128/JB.186.19.6477-6484.2004

Härtig, E., & Jahn, D. (2012). Chapter Five Regulation of the Anaerobic Metabolism in Bacillus subtilis. http://ebookcentral.proquest.com/lib/bu/detail.action?docID=1041274.

Harwood, C. R. ., & Cutting, S. M.. (1990). Molecular biological methods for Bacillus. Wiley.

Hennes, M., Bender, N., Cronenberg, T., Welker, A., & Maier, B. (2023). Collective polarization dynamics in bacterial colonies signify the occurrence of distinct subpopulations. PLoS Biology, 21(1). 10.1371/journal.pbio.3001960

Huang, X., Lao, W., Zhou, Y., Sun, Y., & Wang, Q. (2022). Glutamate dehydrogenase enables *Salmonella* to survive under oxidative stress and escape from clearance in macrophages. FEBS Letters, 596(1), 81–94. 10.1002/1873-3468.14247

Jin, X., Zhang, X., Ding, X., Tian, T., Tseng, C.-K., Luo, X., Chen, X., Lo, C.-J., Leake, M. C., & Bai, F. (2023). Sensitive bacterial V _m_ sensors revealed the excitability of bacterial V _m_ and its role in antibiotic tolerance. Proceedings of the National Academy of Sciences, 120(3). 10.1073/pnas.2208348120

Jones, J. M., & Larkin, J. W. (2021). Toward Bacterial Bioelectric Signal Transduction. Bioelectricity, 3(2), 116–119. 10.1089/bioe.2021.0013

Jones, J. M., Yao, M., Mugler, A., & Larkin, J. W. (2026). Colony morphogenesis regulates sporulation dynamics in bacterial biofilms. Cell Reports, 45(10), 118001. 10.1016/j.celrep.2026.118001

Kaback, H. R. (2015). A chemiosmotic mechanism of symport. Proceedings of the National Academy of Sciences, 112(5), 1259–1264. 10.1073/pnas.1419325112

Kannan, S., Ko, P.-L., Wu, H.-M., & Tung, Y.-C. (2023). Efficient single-cell oxygen consumption rate characterization based on frequency domain fluorescence lifetime imaging microscopy measurement and microfluidic platform. Biomicrofluidics, 17(5). 10.1063/5.0161752

Kirchhoff, C., & Cypionka, H. (2017). Boosted Membrane Potential as Bioenergetic Response to Anoxia in Dinoroseobacter shibae. Frontiers in Microbiology, 8. 10.3389/fmicb.2017.00695

Konkol, M. A., Blair, K. M., & Kearns, D. B. (2013). Plasmid-Encoded ComI Inhibits Competence in the Ancestral 3610 Strain of Bacillus subtilis. Journal of Bacteriology, 195(18), 4085–4093. 10.1128/JB.00696-13

Kralj, J. M., Hochbaum, D. R., Douglass, A. D., & Cohen, A. E. (2011). Electrical Spiking in *Escherichia coli* Probed with a Fluorescent Voltage-Indicating Protein. Science, 333(6040), 345–348. 10.1126/science.1204763

Krasnopeeva, E., Wu, B., Kittler, S., & Guet, C. C. (2026). *Coupling of ATP and PMF in* Escherichia coli *is determined by growth conditions*. 10.64898/2026.04.21.719851

Krüger, L., Herzberg, C., Wicke, D., Bähre, H., Heidemann, J. L., Dickmanns, A., Schmitt, K., Ficner, R., & Stülke, J. (2021). A meet-up of two second messengers: the c-di-AMP receptor DarB controls (p)ppGpp synthesis in Bacillus subtilis. Nature Communications, 12(1). 10.1038/s41467-021-21306-0

Liu, J., Prindle, A., Humphries, J., Gabalda-Sagarra, M., Asally, M., Lee, D. D., Ly, S., Garcia-Ojalvo, J., & Süel, G. M. (2015). Metabolic co-dependence gives rise to collective oscillations within biofilms. Nature, 523(7562), 550–554. 10.1038/nature14660

Lolkema, J. S., Poolman, B., & Konings, W. N. (1998). Bacterial solute uptake and efflux systems. Current Opinion in Microbiology, 1(2), 248–253. 10.1016/S1369-5274(98)80018-0

Martinson, J. N. V., & Walk, S. T. (2020). *Escherichia coli* Residency in the Gut of Healthy Human Adults. EcoSal Plus, 9(1). 10.1128/ecosalplus.esp-0003-2020

Mitchell, P. (1961). Coupling of Phosphorylation to Electron and Hydrogen Transfer by a Chemi-Osmotic type of Mechanism. Nature, 191(4784), 144–148. 10.1038/191144a0

Mulder, O. J., Kostman, M. P., Almodaimegh, A., Edge, M. D., & Larkin, J. W. (2024). *An Agent-Based Model of Metabolic Signaling Oscillations in* Bacillus subtilis *Biofilms*. 10.1101/2024.12.20.629727

Nicholls, D. G. ., & Ferguson, S. J.. (2013). Bioenergetics (4th ed.). Elsevier, Academic Press.

Prindle, A., Liu, J., Asally, M., Ly, S., Garcia-Ojalvo, J., & Süel, G. M. (2015). Ion channels enable electrical communication in bacterial communities. Nature, 527(7576). 10.1038/nature15709

Rocha, R., Teixeira-Duarte, C. M., Jorge, J. M. P., & Morais-Cabral, J. H. (2019). Characterization of the molecular properties of KtrC, a second RCK domain that regulates a Ktr channel in Bacillus subtilis. Journal of Structural Biology, 205(3), 34–43. 10.1016/j.jsb.2019.02.002

Roosild, T. P., Miller, S., Booth, I. R., & Choe, S. (2002). A Mechanism of Regulating Transmembrane Potassium Flux through a Ligand-Mediated Conformational Switch. Cell, 109(6), 781–791. 10.1016/S0092-8674(02)00768-7

Scanlon, T. C., Dostal, S. M., & Griswold, K. E. (2014). A high-throughput screen for antibiotic drug discovery. Biotechnology and Bioengineering, 111(2), 232–243. 10.1002/bit.25019

Schlosser, A., Hamann, A., Bossemeyer, D., Schneider, E., & Bakker, E. P. (1993). NAD ^+^ binding to the *Escherichia coli* K ^+^ -uptake protein TrkA and sequence similarity between TrkA and domains of a family of dehydrogenases suggest a role for NAD ^+^ in bacterial transport. Molecular Microbiology, 9(3), 533–543. 10.1111/j.1365-2958.1993.tb01714.x

Shimoda, L. A., & Polak, J. (2011). Hypoxia. 4. Hypoxia and ion channel function. American Journal of Physiology-Cell Physiology, 300(5), C951–C967. 10.1152/ajpcell.00512.2010

Sierra, J., & Renault, P. (1998). Temporal Pattern of Oxygen Concentration in a Hydromorphic Soil. Soil Science Society of America Journal, 62(5), 1398–1405. 10.2136/sssaj1998.03615995006200050036x

Simoni, R. D., & Postma, P. W. (1975). The Energetics of Bacterial Active Transport. Annual Review of Biochemistry, 44(1), 523–554. 10.1146/annurev.bi.44.070175.002515

Stautz, J., Hellmich, Y., Fuss, M. F., Silberberg, J. M., Devlin, J. R., Stockbridge, R. B., & Hänelt, I. (2021). Molecular Mechanisms for Bacterial Potassium Homeostasis. In Journal of Molecular Biology (Vol. 433, Number 16). Academic Press. 10.1016/j.jmb.2021.166968

Strahl, H., & Hamoen, L. W. (2010). Membrane potential is important for bacterial cell division. Proceedings of the National Academy of Sciences of the United States of America, 107(27), 12281–12286. 10.1073/pnas.1005485107

Stratford, J. P., Edwards, C. L. A., Ghanshyam, M. J., Malyshev, D., Delise, M. A., Hayashi, Y., & Asally, M. (2019). Electrically induced bacterial membrane-potential dynamics correspond to cellular proliferation capacity. Proceedings of the National Academy of Sciences of the United States of America, 116(19), 9552–9557. 10.1073/pnas.1901788116

Tascón, I., Sousa, J. S., Corey, R. A., Mills, D. J., Griwatz, D., Aumüller, N., Mikusevic, V., Stansfeld, P. J., Vonck, J., & Hänelt, I. (2020). Structural basis of proton-coupled potassium transport in the KUP family. Nature Communications, 11(1), 626. 10.1038/s41467-020-14441-7

Teixeira-Duarte, C. M., Fonseca, F., & Morais-Cabral, J. H. (2019). Activation of a nucleotide-dependent RCK domain requires binding of a cation cofactor to a conserved site. eLife, 8. 10.7554/eLife.50661

Tolner, B., Ubbink-Kok, T., Poolman, B., & Konings, W. N. (1995). Characterization of the Proton/Glutamate Symport Protein of Bacillus subtilis and Its Functional Expression in Escherichia coli. In JOURNAL OF BACTERIOLOGY (Vol. 177, Number 10).

Trapp, S., & Ballanyi, K. (1995). KATP channel mediation of anoxia-induced outward current in rat dorsal vagal neurons in vitro. The Journal of Physiology, 487(1), 37–50. 10.1113/jphysiol.1995.sp020859

Yin, W., Cai, X., Ma, H., Zhu, L., Zhang, Y., Chou, S. H., Galperin, M. Y., & He, J. (2020). A decade of research on the second messenger c-di-AMP. In FEMS Microbiology Reviews (Vol. 44, Number 6, pp. 701–724). Oxford University Press. 10.1093/femsre/fuaa019

Young, M., Artsatbanov, V., Beller, H. R., Chandra, G., Chater, K. F., Dover, L. G., Goh, E.-B., Kahan, T., Kaprelyants, A. S., Kyrpides, N., Lapidus, A., Lowry, S. R., Lykidis, A., Mahillon, J., Markowitz, V., Mavromatis, K., Mukamolova, G. V., Oren, A., Rokem, J. S., … Greenblatt, C. L. (2010). Genome Sequence of the Fleming Strain of *Micrococcus luteus*, a Simple Free-Living Actinobacterium. Journal of Bacteriology, 192(3), 841–860. 10.1128/JB.01254-09

Zhou, X., Zhang, N., Xia, L., Li, Q., Shao, J., Shen, Q., & Zhang, R. (2018). ResDE Two-Component Regulatory System Mediates Oxygen Limitation-Induced Biofilm Formation by Bacillus amyloliquefaciens SQR9. Applied and Environmental Microbiology, 84(8). 10.1128/AEM.02744-17

Zong, W., Friedman, E. S., Allu, S. R., Firrman, J., Tu, V., Daniel, S. G., Bittinger, K., Liu, L., Vinogradov, S. A., & Wu, G. D. (2024). Disruption of intestinal oxygen balance in acute colitis alters the gut microbiome. Gut Microbes, 16(1). 10.1080/19490976.2024.2361493

