## Supplementary Information for "Bacteria hyperpolarize in response to oxygen drop"

### Supplementary Information: Modeling Details

#### S1. Model overview

The model was developed to test whether local competition for oxygen can explain the dependence of oxygen-shift-induced hyperpolarization on cell density and ambient oxygen availability. The model explicitly represents individual cells in two dimensions and contains four steps: (i) construction of colony geometry, (ii) definition of the steady-state oxygen field and cell-boundary uptake condition, (iii) self-consistent solution of the oxygen uptake associated with each cell, and (iv) mapping of cell-specific uptake onto a normalized hyperpolarization response. The resulting population response is compared with the experimentally measured mean change in fluorescence intensity over cell-covered regions.

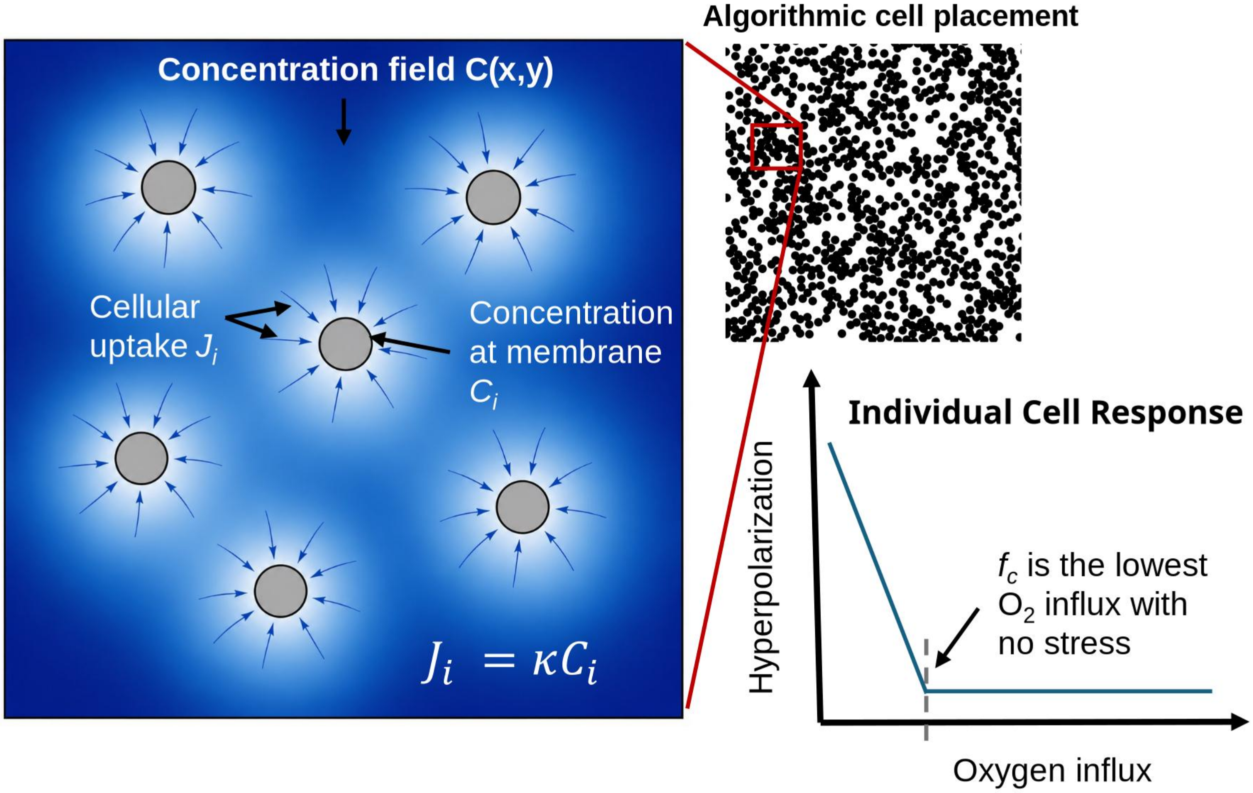


**Figure S1. Overview of the cell-resolved oxygen-competition model.** Cells are placed algorithmically, the extracellular oxygen field is represented by a steady-state diffusion model, oxygen uptake at each cell boundary obeys a Robin condition, and the resulting cell-specific uptake is mapped onto a normalized hyperpolarization response using a critical uptake threshold $f_c$.

#### S2. Colony geometry and simulated cell density

Cells were represented as non-overlapping disks with effective radius a = 1 μm. Cell centers were generated using a modified hard-core Poisson-disk placement procedure with a minimum center-to-center separation of 2a. During placement, a newly initialized cell joined an existing local cluster with probability 0.60; otherwise, it was placed at a random admissible location. For clustering events in the current implementation, an existing cell was selected as a parent and the candidate cell was placed at a random orientation and a center-to-center distance between 2a and 4a, subject to the same non-overlap constraint. This procedure produces heterogeneous cell clusters that qualitatively resemble the spatial organization observed in experimental images.

For model–experiment comparison, simulated cell coverages were matched to the experimentally measured percentage of the field of view occupied by cells (fFOV). To reduce boundary artifacts caused by the absence of cells outside the simulated field, only cells whose centers were located at least one interaction radius R from the domain boundary were included in population-level response calculations. Final results were produced with a square simulation domain of (200+2R)*(200+2R)μm.

#### S3. Steady-state oxygen field and cellular uptake

##### S3.1 Steady-state equation

Because the model describes the steady-state extracellular oxygen field, oxygen concentration C(x,y) satisfies the two-dimensional steady-state diffusion equation in the region outside the cells:

$$D\nabla^2 C = 0 \tag{S1}$$

where D is the effective oxygen diffusion coefficient. The current implementation uses D = 2000 μm² s⁻¹, which is of the order expected for molecular oxygen in aqueous media and dilute gels (St-Denis & Fell, 1971; Bhunia et al., 2016).

##### S3.2 Robin boundary condition

Oxygen uptake at each cell boundary is represented by a Robin (radiation) boundary condition (Collins & Kimball, 1949), in which the uptake flux density is proportional to the local oxygen concentration at the cell surface:

$$D\frac{\partial C_i}{\partial r}|_{r=a} = \kappa C_i(a) \tag{S2}$$

Here, $C_i(a)$ is the oxygen concentration at the boundary of cell *i* and κ is an effective uptake coefficient with dimensions of velocity. Under the sign convention used in the implementation, positive uptake corresponds to an inward oxygen flux. Defining the two-dimensional cell-specific uptake quantity $q_i$ by multiplying the boundary flux density by the disk circumference gives

$$q_i = 2\pi a J_i = 2\pi a\kappa C_i(a) \tag{S3}$$

#### S4. Green-function representation and self-consistent uptake solution

Rather than solving the diffusion equation on a spatial mesh for every colony realization, the extracellular field was represented by superposition of cell-centered Green-function contributions. The signed two-dimensional interaction kernel used in the implementation is

$$G(r) = \{\frac{1}{2\pi D} ln\left(\frac{r}{R}\right), r < R, 0, r \geq R \tag{S4}$$

where r is the center-to-center distance from the source cell and R is an interaction cutoff. For r < R, ln(r/R) < 0, so a positive sink strength decreases the local concentration. The kernel is set to zero at and beyond R.

For cell i, the concentration at its boundary is written as the sum of the background oxygen level and the contributions from all cell sinks:

$$C_i(a) = C_\infty + q_i G(a) + \sum_{j\neq i} q_j G(r_{ij}) \tag{S5}$$

Combining Eq. S5 with the Robin boundary condition in Eq. S3 gives

$$\frac{q_i}{2\pi a\kappa} = C_\infty + q_i G(a) + \sum_{j\neq i} q_j G(r_{ij}) \tag{S6}$$

and therefore the linear system solved in the simulations can be written as

$$C_\infty = q_i\left[\frac{1}{2\pi a\kappa} - G(a)\right] - \sum_{j\neq i} q_j G(r_{ij}) \tag{S7}$$

Thus, for the matrix equation A q = b, the diagonal matrix elements are $A_{ii} = 1/(2\pi a\kappa) - G(a)$, the off-diagonal elements are $A_{ij} = -G(r_{ij})$, and $b_i = C\infty$. Solving this N-cell linear system yields a cell-specific uptake quantity $q_i$ for every simulated cell.

The Green-function interaction was truncated beyond a center-to-center distance R to limit the number and magnitude of contributions from distant cells. The current reported simulations use R = 75 µm. R is a model interaction cutoff and is not interpreted as a physical outer boundary or as a directly measured oxygen diffusion length. A coarse sensitivity study was run over R = [50, 75, 100] and found that errors asymptotically approached a minimum. R = 75 was chosen as a workable compromise between computing time and accuracy.

#### S5. Mapping oxygen uptake to hyperpolarization

The model assumes that cells hyperpolarize when oxygen uptake falls below a critical cell-specific uptake threshold $f_c$. A fixed threshold could not reproduce all four oxygen conditions, so the threshold was allowed to increase linearly with the imposed oxygen level:

$$f_c = f_0 + \alpha * O_{env} \quad \text{(S8)}$$

where $O_{env}$ is the oxygen level after the shift, in percent.

The normalized response $H_i$ of cell i is defined as

$$H_i = \{1 - \frac{q_i}{f_c}, q_i < f_c, 0, q_i \geq f_c \quad \text{(S9)}$$

The population-level simulated response is the mean response over the analyzed cells,

$$\underline{H} = \frac{1}{N_a} \sum_{i=1}^{N_a} \quad H_i \quad \text{(S10)}$$

where $N_a$ is the number of cells in the interior analysis region. This averaging is the model analogue of the experimental analysis, in which the background is masked and the mean fluorescence intensity is calculated over pixels corresponding to cells. The experimental response is the change in this mean intensity before and after the oxygen shift.

#### S6. Model–experiment comparison and parameter fitting

The model was evaluated across the four experimental oxygen conditions (1%, 2%, 5%, and 10% $O_2$) and across simulated coverages corresponding to the experimentally measured fFOV values. A single parameter set ($\kappa$, $f_0$, $\alpha$) was used across all oxygen conditions. For each parameter set, simulated population responses were compared with the corresponding experimentally measured ΔThT for each field of view (Fig. 2e). The measured ΔThT was normalized by a fixed value of 1650 AU, approximately the 1% $O_2$ linear regression (Fig. 2e) extrapolated to full coverage.

In the current fitting implementation, the mean error is calculated as the mean of the root-mean-square errors (RMSEs) for the four oxygen conditions:

$$E_{mean} = (RMSE_{1\%} + RMSE_{2\%} + RMSE_{5\%} + RMSE_{10\%})/4 \quad \text{(S11)}$$

The selected parameter set is the point with the lowest overall model–experiment error, subject to the same geometry-generation and response calculation procedure for all conditions. Parameter sets giving negative uptake for any cell were excluded as unphysical. This occurs for κ above roughly 4000 µm $s^{-1}$.

#### S7. Parameters used in the current model

| Parameter | Description | Value | Units | Comment |
|---|---|---|---|---|
| a | Effective cell-disk radius | 1 | µm | Approximation for *Bacillus subtilis*. |
| D | Oxygen diffusion coefficient | 2000 | µm² $s^{-1}$ | Approximation from $O_2$ in water. |

| | | | | |
|---|---|---|---|---|
| R | Center-to-center interaction cutoff | 75 | µm | Fixed in the reported simulations. |
| P_cluster | Probability of cluster placement | 0.6 | — | Fixed in the reported simulations. |
| $O_{env}$ | Imposed gas-phase oxygen level | 1, 2, 5, 10 | % | Experimental conditions. |
| κ | Effective uptake coefficient | 19.5 | µm $s^{-1}$ | Found with parameter sweep. |
| $f_0$ | Critical two-dimensional uptake threshold, intercept | 4e-19 | same units as $q_i$ | Found with parameter sweep; depends on C∞ scale. |
| α | Critical two-dimensional uptake threshold, slope | 1.66e-19 | Units of $q_i$ per % $O_2$ | Found with parameter sweep. |
| L_box | Simulation domain size | 200 + 2*R | µm | Required for reproducibility. |
| n_rep | Independent colony realizations per condition | 1* | — | Required for reproducibility. |

*A single colony realization was used per condition. Repeating the fit with five independent realizations gave the same best-fit parameter set in every case. Variation between realizations scales as 1/√N, where N is the number of analyzed cells.

#### S8. Parameter-error landscape

The selected parameter set is approximately κ ≈ 20 µm $s^{-1}$ and $f_c$ ≈$4*10^{-19}$+$1.66*10^{-19}$ *$O_{env}$ in the uptake units used by the 2D model. Figure S2 shows two slices through the three-parameter error landscape (κ, $f_0$, α), each passing through the selected parameter set. The κ- $f_0$ slice (Fig. S2 right panel) indicates partial compensation between the effective uptake coefficient and the critical uptake threshold: increasing cellular uptake capacity can be offset, to some extent, by a corresponding change in the uptake threshold used to map oxygen limitation onto hyperpolarization. This compensation is one-sided: values of κ below about 5–10 µm $s^{-1}$ give high error, whereas above this the error remains low up to the negative-uptake limit. The $f_0$–α slice (Fig. S2 left panel) shows a diagonal valley, as $f_0$ and α trade off, with error rising toward α = 0 (an oxygen-independent threshold).

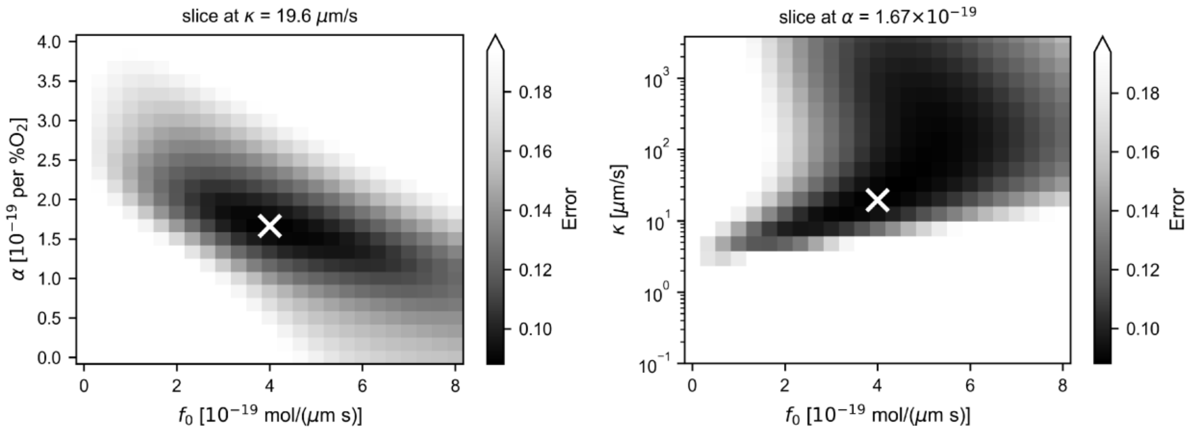


Figure S2. Parameter-error landscape used to identify effective model parameters. Model-experiment error is shown across the scanned values of the uptake coefficient κ, critical uptake threshold intercept and slope $f_0$ and α. The white mark indicates the selected parameter set.

Neither κ nor the threshold parameters $f_0$ and α should be interpreted as uniquely determined physiological constants. They are effective parameters of the reduced two-dimensional model, and their fitted values depend on factors such as the oxygen concentration scale and the mapping between simulated coverage and experimental fFOV. The robust model outputs are therefore the trends across oxygen availability, cell density and spatial arrangement.

#### References


Bhunia, K., Sablani, S. S., Tang, J., & Rasco, B. (2016). Non-invasive measurement of oxygen diffusion in model foods. *Food Research International*, *89*(Pt 1), 161–168.

Collins, F. C., & Kimball, G. E. (1949). Diffusion-controlled reaction rates. *Journal of Colloid Science*, *4*(4), 425–437.

St-Denis, C. E., & Fell, C. J. D. (1971). Diffusivity of oxygen in water. *The Canadian Journal of Chemical Engineering*, *49*(6), 885.
